# Better together against genetic effect heterogeneity: a sex-combined interaction analysis of testosterone levels in the UK Biobank data

**DOI:** 10.1101/2022.06.08.495202

**Authors:** Boxi Lin, Lei Sun

## Abstract

The effect of a genetic variant on a complex trait may differ between female and male, and in the presence of such genetic effect heterogeneity, sex-stratified analysis is often used. For example, genetic effects are sex-specific for testosterone levels, and sex-stratified analysis of testosterone in literature provided easy-to-interpret, sex-specific effect size estimates. However, from the perspective of association testing power, sex-stratified analysis may not be the best approach. As sex-specific genetic effect implies SNP×Sex interaction effect, jointly testing SNP main and SNP×Sex interaction effects may be more powerful than sex-stratified analysis or the standard main-effect testing approach. Moreover, since individual data may be unavailable, it is then of interest to study if the interaction analysis can be derived from sex-stratified summary statistics. We considered several different sex-combined methods and evaluated them through extensive simulation studies. We observed that a) the joint SNP main and SNP×Sex interaction analysis is most robust to a wide range of genetic models, and b) this joint interaction testing result can be obtained by *quadratically* combining sex-stratified summary statistics (i.e. *squared* sum of the sex-stratified summary statistics). We then reanalysed the testosterone levels of the UK Biobank data using sex-combined interaction analysis, which identified 27 new loci that were missed by the sex-stratified approach and the standard sex-combined analysis. Finally, we provide supporting association evidence for nine new loci, uniquely identified by the sex-combined interaction analysis, from earlier association studies of either testosterone level or steroid biosynthesis pathway where testosterone is synthesized. We thus recommend sex-combined interaction analysis, particularly for traits with known sex differences, for most powerful association testing, then followed by sex-stratified analysis for effect size estimation and interpretation.

## 1 INTRODUCTION

The effect of a genetic variant on a complex trait may differ between female and male (Döring et al., 2008; Kong et al., 2008; Sinnott-Armstrong et al., 2021). For example, genetic effects are often sex-specific for hormone-related traits, such as testosterone expression (Sinnott-Armstrong et al., 2021), and severe disease of SARS-CoV-2 infection is known to display sexual dimorphism (Peckham et al., 2020).

A recent sex-stratified genome-wide association study (GWAS) of testosterone levels in the UK Biobank data (Bycroft et al., 2018) identified 79 and 127 independent genome-wide significant signals, respectively, in female and male (Sinnott-Armstrong et al., 2021). Not surprisingly, the top GWAS hits in male were largely non-overlapping with those from female, in terms of both genomic locations and the signaling pathways where the annotated genes are involved. Concerning the emerging GWAS of SARS-CoV-2 susceptibility and hospitalization, although most published work did not perform sex-stratified analysis (or consider SNP×Sex interaction), Roberts et al. (2020) conducted a sex-stratified analysis (followed by a meta-analysis) and identified three genome-wide significant loci in at least one study.

Some earlier GWASs have also reported the existence of genetic effect heterogeneity between female and male. An example for genetic effect in the same direction but different magnitude is the serum uric acid concentrations (Döring et al., 2008): The association between rs7442295 in *SLC2A9* was stronger in female (*p* = 2.6 × 10^−74^, explaining 5.8% of the phenotypic variance) than male (*p* = 7.0 × 10^−17^, explaining 1.2% of the phenotypic variance). There are also examples of genetic effects in opposite directions. For example, the association between recombination rates and SNPs in *RNF212* were reported to be in opposite direction between female and male (Kong et al., 2008).

In the presence of genetic effect heterogeneity between female and male, a sex-stratified analysis approach is often preferred as in Sinnott-Armstrong et al. (2021), which provides easy-to-interpret, sex-specific *effect size estimates* (Li et al., 2010; Anttila et al., 2013; Traylor et al., 2015). However, from the viewpoint of maximizing *power of association testing*, sex-stratified analysis may not be the best analytical strategy for two reasons.

First, after a sex-stratified analysis, it is still tempting to consider sex-combined analysis by, for example, aggregating the association evidence from female and male using the traditional meta-analysis (Cochran, 1954; DerSimonian and Laird, 1986; Willer et al., 2010). However, the fixed-effect meta-approach, though could be used mechanically here, is conceptually at odds with the heterogeneity setting considered here (Greco et al., 2013). The application of the random-effect meta-approach is also challenging, as its implementation requires the between-study variance estimate. When the number of studies (or groups) is small, the estimate is not reliable, resulting in increased type I error rate (Borenstein et al., 2021; Friede et al., 2016).

Second, if the effect of a SNP indeed differs between female and male, it implies SNP×Sex interaction effect. This suggests that an interaction testing approach could be more powerful, but it requires investigation across different genetic models. Additionally, the traditional regression-based mega-approach requires the availability of individual data to include the SNP, Sex and SNP×Sex terms, along with other covariates in the regression model; the corresponding joint SNP main and SNP×Sex interaction 2 degrees of freedom (df) analysis will be termed as the interaction analysis, unless specified otherwise. Thus, it is another intriguing question if the 2 dfinteraction analysis can be derived from sex-stratified summary statistics.

To answer these questions and compare powers between sex-stratified and different sex-combined approaches, let *Z_F_* and *Z_M_* be the summary statistics from the sex-stratified analysis, and *p_F_* and *p_M_* the corresponding p-values, respectively for female and male. Now consider three different sex-combined approaches: (for completeness) the classical meta-analysis of using a weighted average of *Z_F_* and *Z_M_* (*TSG_L_*), the minimal p-value of *p_F_* and *p_M_* (*TSG_min_*), and an omnibus test of using a weighted sum of *squared Z_F_* and *Z_M_* (*TSG_Q_*, 2 df) (Li and Lagakos, 2006; Derkach et al., 2014). For the mega-analysis of individual data, if available, let *T I_joint_* be the 2 df test derived from jointly testing both the main and SNP×Sex interaction effects via regression. For completeness, let’s also consider the most commonly used GWAS approach (Sullivan et al., 2013), testing the genetic main effect (*T_main_*) in a regression model without the SNP×Sex interaction term, but with sex included as a covariate.

Without loss of generality, assume the genetic effect in female (*b_F_*) is always greater or equal to that in male (*b_M_*), where *b_M_* can be zero or in a different direction from *b_F_*. If *b_M_* = 0, intuitively, the female-only *TSG_F_* should be the most powerful test. Similarly, if *b_M_* = *b_F_*, then *T_main_* should be the most powerful test. However, in both cases, it is unclear if the 2 df *T I_joint_* test would be competitive, and if the 2 df *TSG_Q_* behaves similarly. Such an understanding is important for recommending a test that performs reasonably well across different scenarios.

Focusing on gene×environmental interaction analysis of a binary trait, the important work of Kraft et al. (2007) concluded that “Although the joint test of genetic marginal effect and interaction is not the most powerful, it is nearly optimal across all penetrance models we considered.” (Note that sex can be statistically treated as an environmental covariate.) Additionally, in a different study setting where Chen et al. (2021) compared the 2 df genotypic test with the 1 df additive test, derived from a correctly specified 1 df additive model, the maximum power loss of the 2 df test was theoretically capped at 11.4% across all parameter values and sample sizes; this is because “the derivation is based on ncp [non-centrality parameter] and *α* alone.” Thus, we can deduce that *T I_joint_* is a more robust test than *T_main_*,

However, insights are lacking on the method comparison between *T I_joint_* and the other tests, including *TSG_F_*, *TSG_min_*, *TSG_L_*, and *TSG_Q_*. The relative performance between (some of) these tests have been compared through multiple other empirical studies (Magi et al., 2010; Aschard et al., 2010; Clarke and Morris, 2010; Sung et al., 2016, 2014), and the perspective piece by Aschard (2016) offers great insights into the “contribution of interaction effects to the variance of quantitative outcomes”. But these earlier studies focused on the testing of the gene×environment interaction term, rather than studying if the potential heterogeneous genetic effects can be *leveraged* through gene×environment (here SNP×Sex) interaction analysis, which is the main objective of our work here.

In the remainder of this report, first, we describe the six association testing methods considered and the simulation study design for method evaluation. Second, we present the simulation results, comparing the performance of the six tests across different scenarios, with varying degrees of genetic effect heterogeneity and male-to-female sample size ratio. We show that a) there is no uniformly most powerful test across all scenarios, b) *T I_joint_* and *TSG_Q_* are more robust than the others, and c) *T I_joint_* and *TSG_Q_* are practically the same for interaction analysis, complementing the known result that *T_main_* and *TSG_L_* are asymptotically equivalent for studying genetic main effect (Lin and Zeng, 2010). Third, we revisited the sex-stratified GWAS of testosterone levels in the UK Biobank data by Sinnott-Armstrong et al. (2021). After first replicating their sex-stratified findings, we then show that sex-combined *interaction* analysis, *T I_joint_* and *TSG_Q_* (if only summary statistics are available), are indeed more powerful than sex-stratified GWAS and the traditional sex-combined analysis (*T_main_*, *TSG_L_* or *TSG_min_*), in terms of both the number and strength of the signals. Finally, we replicated some of the new findings using the GWAS catalogue resource (Buniello et al., 2019). We conclude the report with additional remarks and a discussion of future work.

## 2 MATERIALS AND METHOD

### 2.1 Notation and set-up

Let *n_F_* and *n_M_* = *k* · *n_F_* be the sample sizes of, respectively, the female and male samples, where *k* is the male-to-female sample size ratio. Let *n* = *n_M_* + *n_F_* be the total sample size of the combined sample. Let 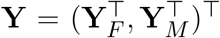 be the phenotype vector of a trait of interest, 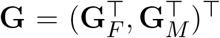 be the genotypes of a SNP under the study, and **S** be the *n* × 1 sex-indicator vector. For each element in **S**, without loss of generality, we define *S_i_* = 1 for a female, *i* = 1, …, *n_F_*, and *S_j_* = 0 for a male, *j* = 1, …, *n_M_*. We assume the phenotype data were generated from the following generalized linear model,

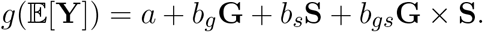

In this work, we mainly focus on the quantitative trait with identity link, but the analysis is generalizable to other types of trait with appropriate link functions.

### 2.2 *TSG_F_* and *TSG_M_*, the female and male sex-stratified analysis

As a SNP’s genetic effect could be sex-specific, we first consider the commonly used sex-stratified approach to testing association between **Y** and **G**. In practice, important environmental covariates **Z**’s are included in the analysis, but they are omitted here for notation simplicity but without loss of generality; results are characteristically the same unless (beyond gene×sex) gene×environmental interaction analysis is also of interest, which is beyond the scope of this work.

Let

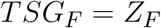

be the standard Wald test statistic, derived from the regression model, 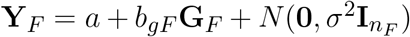, by testing *H*_0_ : *b_gF_* = 0. (If the trait is binary, a logistic regression will be used.) *Z_F_* follows *N*(0, 1) asymptotically under *H*_0_, and the asymptotic p-value of *TSG_F_* is *p_F_* = 2 × (1 − Φ[|*Z_F_*|]), where Φ is the cumulative density function of the standard normal distribution. Similarly, we can obtain *Z_M_* and *p_M_* for the association analysis using the male sample.

### 2.3 Three sex-combined approaches based on sex-stratified summary statistics

#### 2.3.1 *TSG_L_*, the classical meta-analysis

As discussed earlier, the traditional fixed-effect meta-analysis is not recommended for the heterogeneity setting considered here, and the random-effect approach is not applicable to studies of two groups only (Borenstein et al., 2021). But for completeness of our evaluation, we include *TSG_L_*, the classic inverse-variance weighted average of *Z_F_* and *Z_M_*,

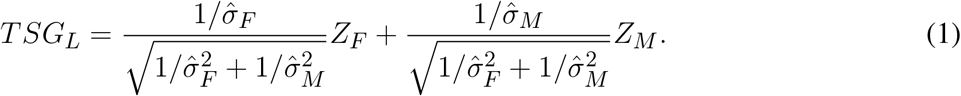

Under the null of no association, *TSG_L_* is asymptotically *N*(0, 1) distributed.

The subscript _*L*_ reflects the fact that *TSG_L_* combines the two *Z* statistics linearly. As a result, *TSG_L_* could lose power when the genetic effects in female and male have opposite directions. This is akin to the burden type of tests for joint analyzing multiple rare variants (Derkach et al., 2014).

#### 2.3.2 *TSG_Q_* (2 df), an omnibus test

Instead of forming a directional test by linearly combining *Z_F_* and *Z_M_*, we could construct an omnibus test by calculating a quadratic sum as

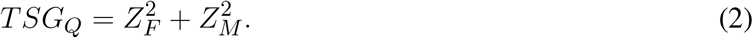

Under the null of no association, *TSG_Q_* asymptotically follows the centralized 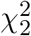 distribution. A sample-size-based weighted sum of 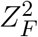 and 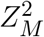 can be also considered, but does not alter the results qualitatively (results not shown). Additionally, *TSG_Q_* appears to be numerically connected with the interaction analysis, which we discuss in Section 2.4.1.

#### 2.3.3 *TSG_min_*, the minimal p-value approach

We could also consider the minimal p-value approach,

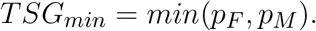

Note that *p_M_* and *p_F_* are independent of each other, and each is Unif(0, 1) distributed under the null of no association. Thus, the cumulative distribution function of *TSG_min_* is *F*(x) = 1 − (1 − *x*)^2^, which is used to obtain p-value for *TSG_min_*-based association analysis.

### 2.4 Two sex-combined mega-analysis approaches

When individual data are available, we could consider the following two mega-analysis approaches through regression, with or without the SNP×Sex interaction term; Sex is always included as a covariate by convention.

#### 2.4.1 *T I_joint_*, the 2 df main and interaction testing approach

We first consider the following regression model,

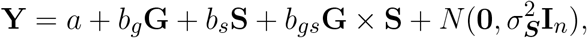

where the interaction term is included to capture the potential genetic heterogeneity between female and male. Additionally and importantly, the error model is sex-specific, 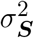. That is, sex-specific variances, 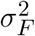 and 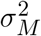, are assigned to the female and male groups, respectively; for example, human height differs between female and male in both its mean and variances (Deng et al., 2019).

We use

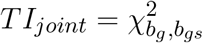

to denote the 2 df Wald test statistic derived from jointly testing *H*_0_ : *b_g_* = *b_gs_* = 0. This test has one extra df than the traditional main-effect-only 1 df test. However this does not directly translate to loss of power, as the corresponding non-centrality parameters may differ (Chen et al., 2021). We provide numerical evidence in Sections 3 and 4.

#### 2.4.2 *T_main_*, the genetic main effect only testing approach (and adjusting for sex main effect)

For completeness, we also consider the standard approach of testing for genetic main effect only. Although such a main-effect only test can be derived from the interaction regression model above, the resulting test is difficult to interpret in the presence of an interaction effect; similarly it is difficult to interpret the interaction effect in the presence of the main effect (Brambor et al., 2006; Aschard, 2016).

Thus, we consider the following most commonly used regression model,

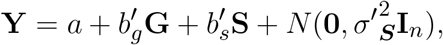

with the modification on the error term to allow for sex-specific variance 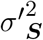, as noted earlier. Additionally, important environmental factors should be included in the regression but omitted here to ease notation. We use

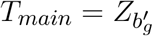

to denote the Wald test statistic derived from testing 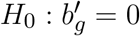.

### 2.5 Simulation study design

We conducted extensive simulations to evaluate the performance of the six association tests, namely *TSG_F_*, *TSG_min_*, *TSG_L_*, *TSG_Q_*, *T_main_*, and *T I_joint_*, where *TSG_min_*, *TSG_L_* and *TSG_Q_* are summary-statistics-based, and *TSG_Q_* and *T I_joint_* are 2 df tests. As genetic effect in male was assumed to be no larger than that in female, *TSG_M_* was not explicitly examined but used to construct the summary statistics-based sex-combined analyses.

The alternative scenarios considered generally follow those in Magi et al. (2010), where genetic effects could be the same between female and male or sex-specific. Specifically, we considered the bi-allelic SNP of interest *G* is coded additively as in convention, and ℙ(*G* = 2) = *maf*^2^, ℙ(*G* = 1) = 2 × *maf* × (1 − *maf*), and ℙ(*G* = 0) = (1 − *maf*)^2^, where maf is the population minor allele frequency and takes values of *maf* = 0.05, 0.1 or 0.25. We also considered varying male-to-female sample size ratio, with *k* = *n_M_*/*n_F_* taking values from {0.5, 1, 1.5, 2}, and *n_F_* = 5, 000. As power depends on both sample size and effect size, we did not empirically consider different *n_F_*, as it does not affect the *relative* performance in power between the different methods. The sex-combined phenotype generating regression model can be reformulated, equivalently, as two separate simulation models for male and female, respectively, as

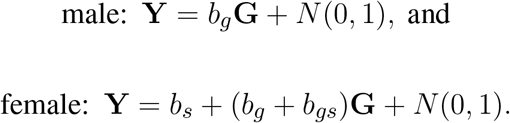

We fixed *b_s_* = 1 so that sex explained about 20% of the total phenotypic variation. By varying *b_g_* and *b_gs_*, we were able to simulate phenotype data under different association scenarios including;

• The null scenario: No genetic effects in both female and male.

To evaluate the empirical type I error rates, we fixed *b_g_* = *b_gs_* = 0 to generate data under the null hypothesis of no genetic effects of *G* on *Y*, in both female and male.

• Alternative scenario A1: Homogeneous genetic effect between female and male.

In scenario A1, we fixed *b_gs_* = 0 so that the true genetic model has no interaction effect, which is statistically equivalent with homogeneous genetic effect between female and male. This is the best-case scenario for *T_main_* and the worst for *T I_joint_*, but we will show that power of *T I_joint_* is competitive under A1, and *TSG_Q_* has the same performance as *T I_joint_*.

The range of *b_g_* was [0.06, 0.3], corresponding to the phenotypic variation explained by the genetic variable 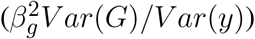 in the range of [0.14%, 3.24%]. For completeness, we also let *b_g_* = 0 so the power study also includes the null scenario, where we expect the empirical power to be at the nominal type I error level. Similar inclusion of the null was made when studying alternative scenarios A2 and A3 below.

• Alternative scenario A2: Female-only genetic effect.

In scenario A2, we fixed *b_g_* = 0 (no effect in male), and *b_gs_* in the range of [0.06, 0.3] (the effect in female). This setting is the best-case scenario for *TSG_F_*. But again, we will show that *T I_joint_* and *TSG_Q_* are competitive, as compared with *TSG_F_*, and under A2 *T I_joint_* and *TSG_Q_* also have the same empirical performance.

• Alternative scenario A3: Heterogeneous genetic effect between female and male.

Here we fixed genetic effect in female at *b_g_* + *b_gs_* = 0.15, and *b_g_* in the range of [−0.6, 0.3] for the genetic effect in male. When *b_g_* < 0, the effect directions differ between female and male. When *b_g_* > 0, although the effect directions are the same, the effect magnitudes differ, unless *b_g_* = 0.15.

Additionally, we conducted a sensitivity analysis under the following two settings;

• Sensitive Study 1: Non-normal residual distributions.

Instead of the standard normal residual, we generated *ϵ* from student-t_4_ and 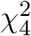 to compare the performance of the testing methods when there are, respectively, excess kurtosis and skewness in the residual distributions.

• Sensitive Study 2: Dominant genetic model.

Instead of the standard additive genetic model, we generated phenotypes from dominant models, which statistically also covers recessive models by switching the choice of the baseline allele. The working models remained additive, following the common GWAS practice.

Finally, although we do not expect the relative method performance to change when analyzing a binary trait, for completeness, we also generated binary phenotype data from *Bernoulli*(*p*), where *p* = *g*^−1^(*b_g_***G** + *b_s_***S** + *b_gs_***G** × **S**) and *g* is the logit link function.

## 3 SIMULATION RESULTS

Results here focus on power across different alternative scenarios, as all six tests are standard statistical tests derived directly or indirectly from regression and expected to be accurate, particularly when the trait is normally distributed. But for completeness, we report the empirical type I error rates under the null of no association and including the sensitivity analyses.

### 3.1 Empirical type I error rates

We used *R* = 5 × 10^6^ replications to evaluate the empirical type I error rates at the nominal level of *α* = 10^−5^. Results in Figure S1 show that, as expected, all six tests are accurate, and have the nominal level *α* = 10^−5^ covered by 95% binomial proportion confidence interval 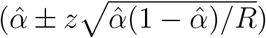, where 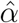 is empirical type I error. We note that *α* = 5 × 10^−8^ (Dudbridge and Gusnanto, 2008) is ideal but requires at least *R* = 10^10^ replicates for accurate type I error evaluation, for each sample size considered, which is computational expensive, but *α* = 5 × 10^−8^ was used for power study.

### 3.2 Power comparison

Figure 1 shows the empirical powers of the six testing methods, respectively, under the three alternative scenarios, A1, A2 and A3, where the empirical powers, at the genome-wide significance level of 5 × 10^−8^, were obtained from 10^5^ replicates. Result presentation here focuses on sample sizes of *n_F_* = *n_M_* = 5, 000. Results for unbalanced sample size ratio with *k* ∈ {0.5, 1.5, 2} are characteristically similar in that the relative method performance stays the same (Figures S2-S4).

**Figure 1.**
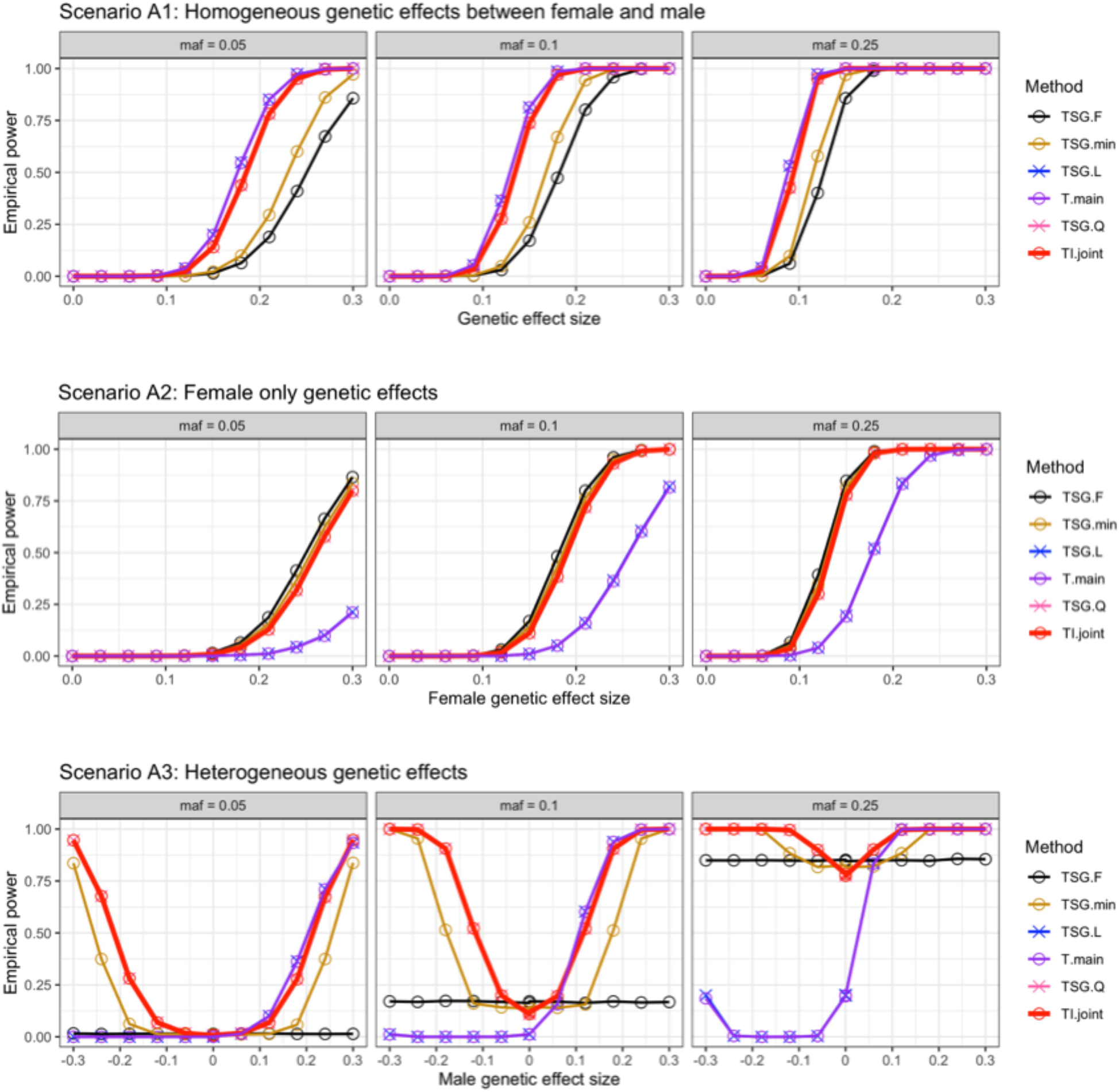
Power comparison at *α* = 5 × 10^−8^ with female sample size *n_F_* = 5, 000 and *k* = 1 (*n_M_* = 5, 000) under the three alternative scenarios, A1, A2 and A3, stratified by the MAF. A1: Homogeneous genetic effect between female and male, and the genetic effect sizes ranged from 0 to 0.3; A2: Female-only genetic effect, and the genetic effect sizes ranged from 0 to 0.3; A3: Heterogeneous genetic effect between female and male, the genetic effect in female was kept at 0.15, while the effect in male ranged from −0.3 to 0.3. Six association testing methods were evaluated: (1) The female-only sex-stratified analysis (*TSG_F_*); (2) The minimum p-value of sex-stratified analysis (*TSG_min_*); (3) A meta-analysis combining Z-scores (*TSG_L_*); (4) The main-effect-only model (*T_main_*); (5) An omnibus test combining squared Z-scores (*TSG_Q_*), the recommended test when only sex-stratified summary statistics are available; (6) The recommended joint test of both the main and interaction effects (*T I_joint_*). Results for other *k* = *n_M_* /*n_F_* male-to-female ratios are shown in Figures S2-S4.

In Scenario A1 where genetic effects are the same between female and male, it is easy to see that this is the best case-scenario for *T_main_*, which jointly analyzes all samples available through a (correctly-specified) main-effect-only regression model. Similarly, *TSG_F_* should have the lowest power, as the female-only analysis ignores the *n_M_* = *k* · *n_F_* of the total available n samples. Additionally, when there is no effect heterogeneity, meta-analysis (linearly combining *Z_F_* and *Z_M_*) is as efficient as mega-analysis (Lin and Zeng, 2010). Thus, *TSG_L_* has the same power as *T_main_* as expected; the two power curves overlap in Figure 1 Scenario A1.

As there is no interaction effect in Scenario A1, the 2 df *T I_joint_* test, jointly testing both the main and interaction effects, is expected to be less powerful than *T_main_*. However, results in Figure 1 Scenario A1 show that the loss of power is marginal, suggesting the robustness of *T I_joint_* even under the scenario of no interaction effect. Additionally, *T I_joint_* is noticeably more powerful than the minimum-p value approach of *TSG_min_*. Finally, the power curve of *T I_joint_*, interestingly, overlaps with that of *TSG_Q_* (quadratically combining 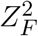 and 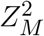.)

In Scenario A2, where the genetic effect exists only in female, the female-only sex-stratified test *TSG_F_* is most powerful as expected, and *T_main_* is the least powerful with significant loss of power; power of *TSG_L_* is practically identical to that of *T_main_* as expected. Compared with Scenario A1, the performances of *TSG_F_* and *T_main_* (and *TSG_L_*) here in Scenario A2 are reversed, suggesting that neither method is robust against different alternatives, which are unknown in practice. Also as expected is *TSG_min_* being almost as powerful as *TSG_F_*, as *TSG_min_* captures the association signal through the female-only sex-stratified analysis; the slight loss of efficiency of *TSG_min_* is a result of searching for the ‘best’ result between the two groups, akin to *p*-hacking penalization.

Importantly, *T I_joint_* is also competitive in this case, with power almost identical with that of *TSG_min_*, and only slightly smaller than that of the most powerful method of *TSG_F_*. Interestingly, same as in Scenario A1, the power curve of *T I_joint_* overlaps with that of *TSG_Q_* in Scenario A2 as well.

In Scenario A3 where genetic effects exist in both female and male but differ in magnitude and/or direction, it reasonable to predict that no method dominates the others as confirmed by results in Figure 1. Specifically, (I) When the genetic effect in male is close to zero, the relative performance of the different methods is, as expected, similar to that observed scenario A2: The female-only sex-stratified test *TSG_F_* has the highest power, but *TSG_min_*, *TSG_Q_* and *T I_joint_* are competitive. (II) When the genetic effect in male is close to 0.15 (i.e. without effect heterogeneity between male and female), the relative performance is similar to that in scenario A1: *T_main_* and *TSG_L_* have the highest power, but *TSG_Q_* and *T I_joint_* are competitive. (III) When the genetic effect in male is close to −0.15 (i.e. with severe effect heterogeneity to the extend of opposite effect directions), *TSG_Q_* and *T I_joint_* are clearly more powerful than the other methods.

Finally, across the range of parameter values under Scenario A3, the empirical powers of *T I_joint_* and *TSG_Q_* are the same as under A1 and A2. Thus, although theoretical justification is needed, our simulation results suggest that the 2 df interaction mega-analysis can be obtained from sex-stratified summary statistics through *TSG_Q_* (quadratically combining 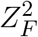 and 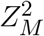). We will explore this as future work from the theoretical lens, following the earlier work of Aschard et al. (2010) and Aschard (2016) that focused on Score tests and regression models with residual variances independent of *E* (Sex in our case).

### 3.3 Sensitivity studies with model mis-specification and binary outcome

When the error distribution is *t*_4_ or 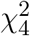, given our simulation sample size (*n_F_* = 5, 000 with male-to-female sample size ratio of 0.5, 1, 1.5, or 2), the empirical type I error rates can be slightly inflated when MAF is low at 0.05, for all six tests examined (Figure S5). This is, however, not surprising as the convergence of the least square estimator’s sampling distribution is slower when the residual distribution is non-normal, particularly asymmetrical (e.g. 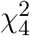) (Lehmann, 1999). Subsequently, we did not consider power evaluation with non-normal residuals.

When the phenotype values are simulated from a dominant genetic model while the working model is additive, or the trait is binary, the type I error control results are similar to above (Figures S6 and S7).

Finally, across the three alternatives considered, the relative performances of the different tests for dominant generating models (Figure S8, the right panel) and for binary traits (Figure S9, left panel for additive model and right panel for dominant model) remain consistent with those observed from normally distributed traits and additive genetic models, as shown Figure 1 and Figures S2-S4.

## 4 APPLICATION TO GWAS OF THE TESTOSTERONE LEVELS IN THE UK BIOBANK DATA

Utilizing the UK Biobank data (Application number: 64875), we conducted GWAS of testosterone levels to compare the performance of sex-stratified and different sex-combined (main only vs. main and interaction joint) testing approaches.

First, we replicated the summary statistics, *Z_F_* (named *TSG_F_* for test of sub-group) *Z_M_* (*TSG_M_*), of the sex-stratified GWAS of Sinnott-Armstrong et al. (2021) in accordance with their sample quality control and association testing procedures. With ~ 320, 000 unrelated white British individuals as the study sample, we conducted GWAS using ~ 800, 000 genotyped SNPs with *MAF* > 0.01; the sample size here is about 32 times larger than that of the simulation study, which allowed us to lower the MAF threshold from 5% to 1%. Our results are consistent with summary statistics provided in Sinnott-Armstrong et al. (2021) (https://figshare.com/collections/Supplementary_Data_For_Sinnott-Armstrong_and_Naqvi_eLife_2020/5304500/1). Second, we calculated summary statistics-based sex-combined test statistics, *TSG_L_* (weighted sum of *Z_F_* and *Z_M_*) and *TSG_Q_* (sum of 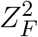 and 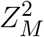), and their corresponding p-values, as described in Section 2; we omitted *TSG_min_* from the application study, as our simulation studies has shown that it is sub-optimal as compared with *TSG_Q_*. Lastly, we conducted the regression analyses using the individual data as described in Section 2.3.3, to obtain *T_main_* and *T I_joint_*.

Similar to the simulation results, here we observed that *TSG_L_* and *T_main_* are empirically equivalent (Figure S10, the right plot), which is expected given the theoretical results in Additionally, *TSG_Q_* and *T I_joint_* (Figure S10, the left plot) are also empirically equivalent, which is also not surprising given the work of Aschard et al. (2010) and Aschard (2016). However, additional theoretical work is needed as the earlier work focused on Score tests and covariate-independent residual variances.

The Miami plots in Figure 2 clearly demonstrate that *TSG_Q_* was able to identify the top hits detected by either *TSG_F_* (Figure 2a) or *TSG_M_* (Figure 2b), as well as identify additional new loci. In contrast, as expected, *TSG_L_* performs less well than *TSG_Q_* in the presence of sex-effect heterogeneity. To provide better visualization for all significant loci, Figures S11 and S12 provide the Miami plots with −*log*_10_(*p*) truncated at 50 and 10, respectively.

**Figure 2.**
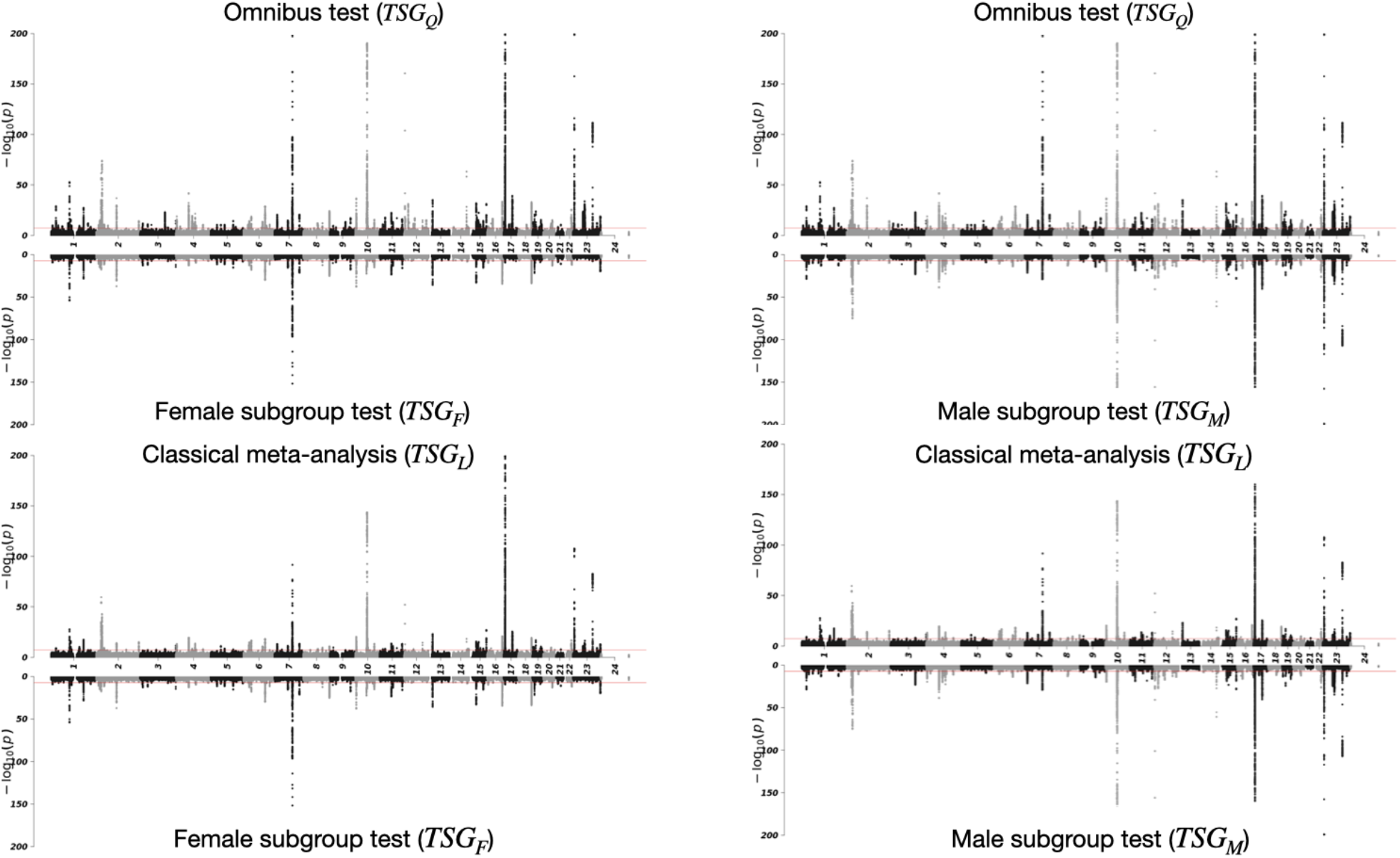
Miami plots comparing the − log_10_ (*p*) between tests based on (a) *TSG_Q_* and *TSG_F_*, (b) *TSG_Q_* and *TSG_M_*, (c) *TSG_L_* and *TSG_F_*, and (d) *TSG_L_* and *TSG_M_*, leveraging the sex-stratified GWAS summary statistics of testosterone level from Sinnott-Armstrong et al. (2021). Here *TSG_F_* and *TSG_M_* are sex-stratified GWAS results of Sinnott-Armstrong et al. (2021). The omnibus *TSG_Q_* results were obtained using equation 2, and the traditional meta-analysis *TSG_L_* results were obtained using equation 1. The red horizontal lines indicate the genome-wide significant threshold of 5 × 10^−8^ on the −*log*_10_ scale. The maximum −*log*_10_ *p* was truncated at 200 for better visualization. Miami plots with −*log*_10_(*p*) truncated at 50 and 10 are shown in Figures S11 and S12, respectively.

To quantitatively evaluate and compare the performance between sex-stratified and sex-combined methods, the Venn diagram in Figure 3 summarizes the counts of independent loci. The independent loci were defined as the LD blocks constructed by Haploview using solid spline method (Barrett et al., 2004), with LD threshold *r*^2^ = 0.1 and window size of 100kb. Figure 3 clearly shows that, unlike *TSG_L_*, *TSG_Q_* was able to retain most of the signals identified from either *TSG_F_* or *TSG_M_*, as well as discover new loci.

**Figure 3.**
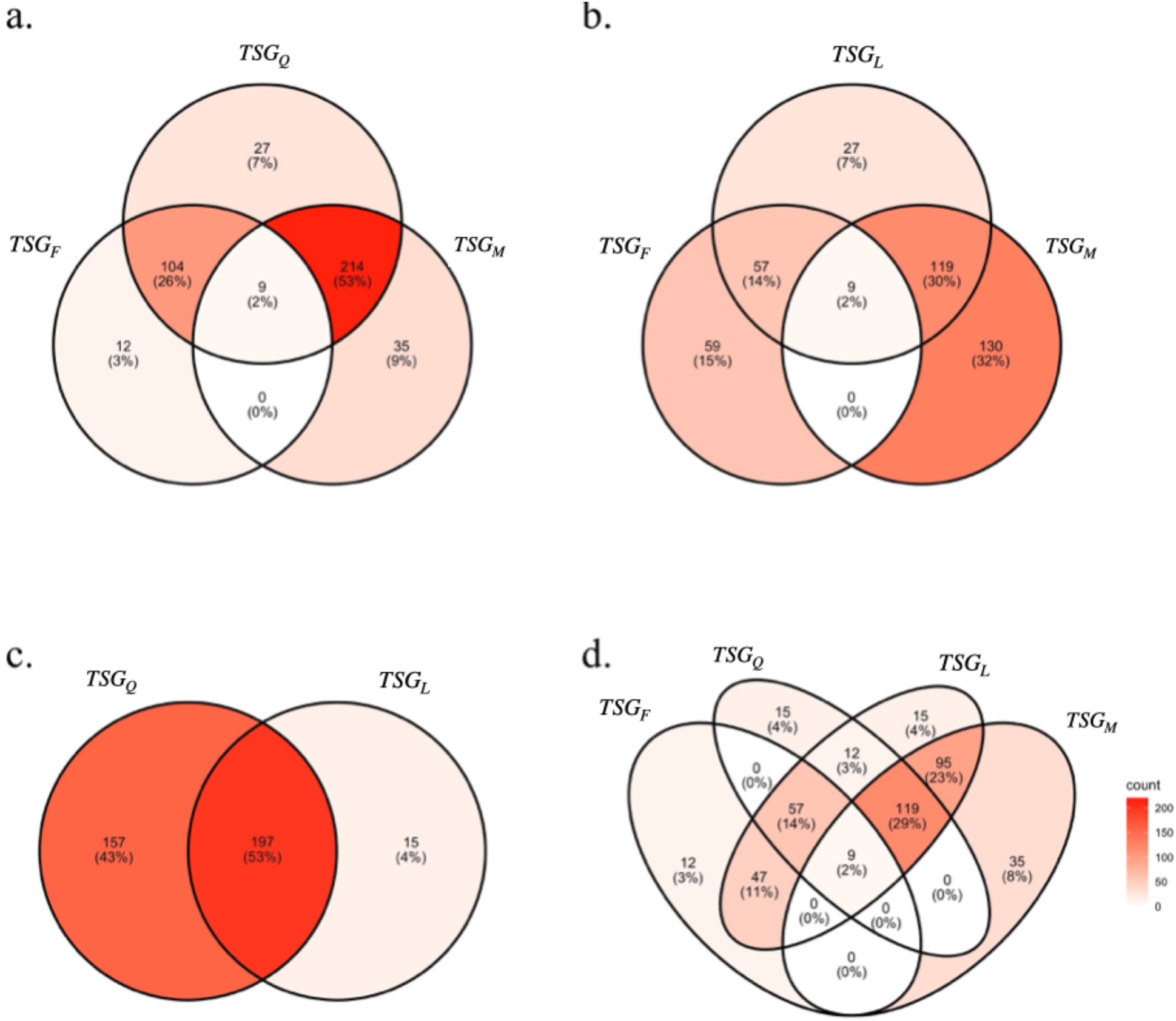
Venn diagrams showing the relationship between the independent genome-wide significant loci identified from (a) *TSG_Q_* and the sex-stratified tests, *TSG_F_* and *TSG_M_*, (b) *TSG_L_* and the sex-stratified tests, (c) *TSG_Q_* and *TSG_L_*, and (d) All of the four tests. Here *TSG_F_* and *TSG_M_* are sex-stratified GWAS results of (2021). The omnibus *TSG_Q_* results were obtained using equation 2, and the traditional meta-analysis *TSG_L_* results were obtained using equation 1. The independent loci were defined as the LD blocks constructed by Haploview using solid spline method (Barrett et al., 2004), with LD threshold *r*^2^ = 0.1 and window size of 100kb.

Additionally, the PP plots in Figure 4 show that for the genome-wide significant SNPs identified by *TSG_Q_* or another method (sex-stratified or *TSG_L_*), visually they all appear in the bottom triangle of each pair-wise comparison. This means that *TSG_Q_* is more robust and can be much more powerful than the other testing methods, as for the associated variants, the − log_10_(*p*) of *TSG_Q_* are always almost as large as or substantially larger than the − log_10_(*p*) of the other tests.

**Figure 4.**
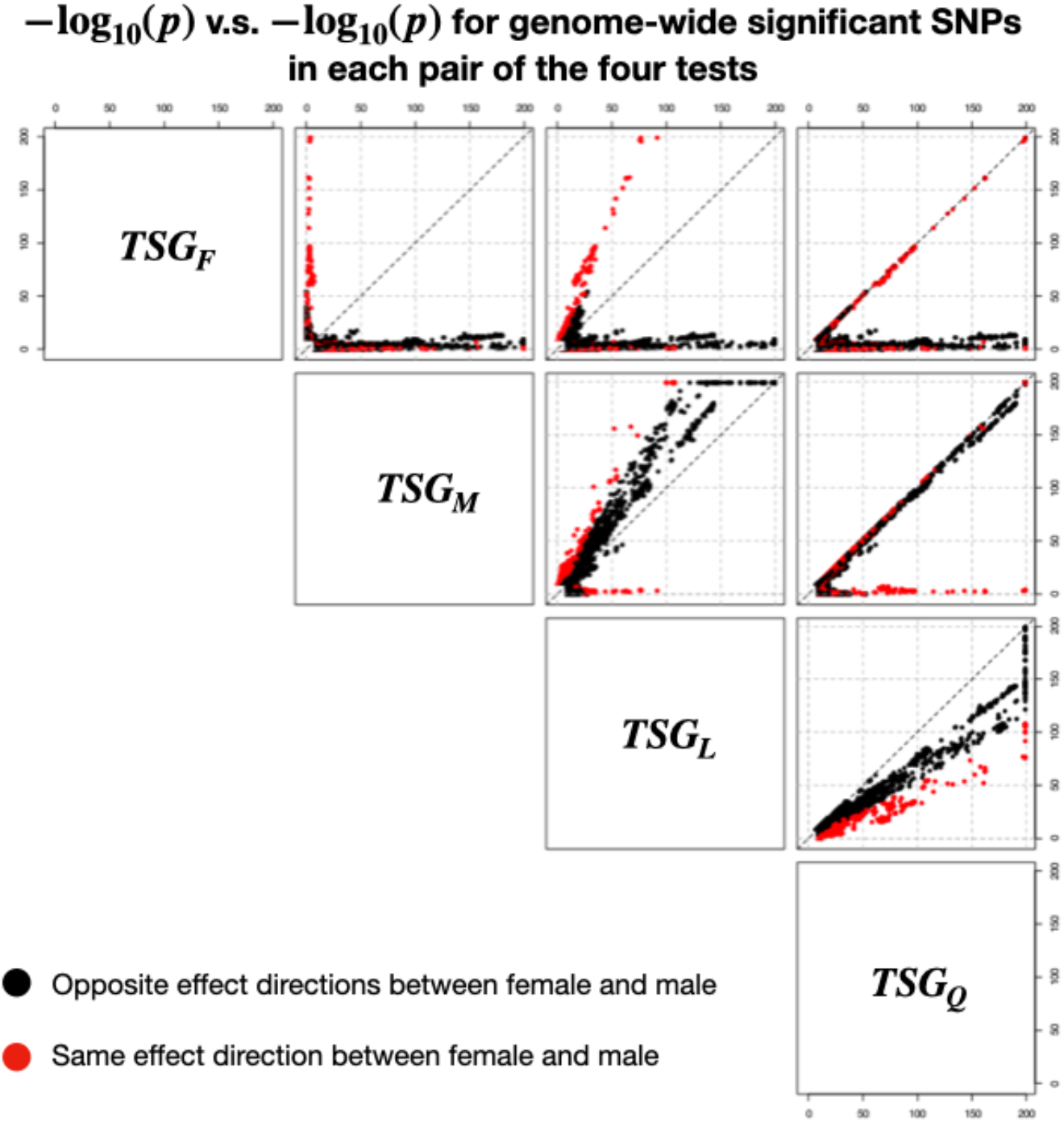
Pairwise PP plots comparing the −log_10_ *p*-values between the female-only sex-stratified analysis (*TSG_F_*), male-only sex-stratified analysis (*TSG_M_*), the classical meta-analysis (*TSG_L_*), and the omnibus test combining squared Z-scores (*TSG_Q_*). Here *TSG_F_* and *TSG_M_* are sex-stratified GWAS results of (2021). The omnibus *TSG_Q_* results were obtained using equation 2, and the traditional meta-analysis *TSG_L_* results were obtained using equation 1. In each sub-plot, each point represents a SNP reaching genome-wide significant level of 5 × 10^−8^ *in any of the two corresponding testing methods*. The maximum − log_10_ *p* was truncated at 200 for better visualization. PP plots with −*log*_10_(p) truncated at 50 and 10 are shown in Figures S13 and S14, respectively.

Indeed, although *TSG_Q_* missed 12 and 35 independent loci identified, respectively, from the subgroup analyses *TSG_M_* and *TSG_F_* (Figure 3a), the PP plots in Figures 5c and 5d show that for these SNPs the maximum − log_10_(*p*) of *TSG_F_* and *TSG_M_* was capped at 8.17, while the minimum − log_10_(*p*) of *TSG_Q_* was bounded at 7.30. That is, even *TSG_Q_* may miss some signals strictly by the genome-wide significant threshold, the testing results of *TSG_Q_* are comparable with that *TSG_F_* and *TSG_M_*, in terms of ranking of the associated SNPs. In contrast, the PP plots in Figures 5a and 5b show that among the novel signals identified by *TSG_Q_*, the − log_10_(*p*) of *TSG_Q_* could be as high as 12.52 while the − log_10_(*p*) of sex-stratified tests could be as low as 1.36, suggesting that *TSG_Q_* is much more powerful than sex-stratified tests.

**Figure 5.**
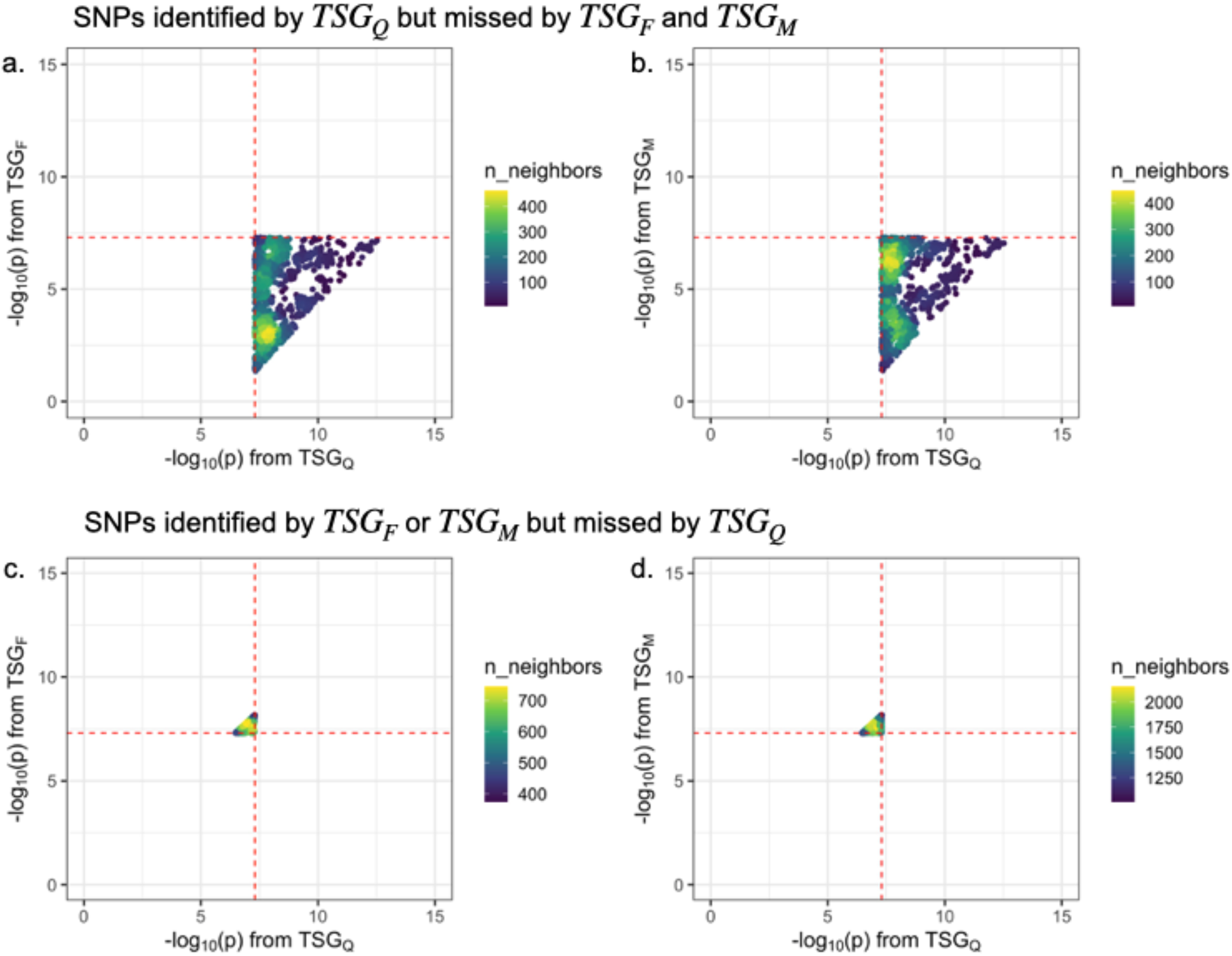
Restricted pairwise PP plots, considering (a) and (b): SNPs identified by *TSG_Q_* but missed by *TSG_F_* and *TSG_M_*, (c) and (d): SNPs identified by *TSG_F_* or *TSG_M_* but missed by *TSG_Q_*. Here *TSG_F_* and *TSG_M_* are sex-stratified GWAS results of Sinnott-Armstrong et al. (2021). The omnibus *TSG_Q_* results were obtained using equation 2, and the traditional meta-analysis *TSG_L_* results were obtained using equation 1. The red lines indicate the genome-wide significant threshold of 5 × 10^−8^ on the −*log*_10_ scale. The color of points indicates *n_neighbors*, i.e. the number of its neighbouring points of the point. The p-values were *not* truncated for the set of SNPs considered.

Finally, we evaluated whether the 27 novel genome-wide significant loci, uniquely identified by *TSG_Q_* but missed by *TSG_F_*, *TSG_M_* and *TSG_L_* in the UK Biobank data, are biologically relevant to the testosterone level phenotype. According to the GWAS catalog (Buniello et al., 2018), nine out of the 27 loci were previously reported to be associated with either testosterone level or steroid biosynthesis pathway where testosterone is synthesized (Haas et al. (2022); Ruth et al. (2020);Richardson et al. (2022);Hoffmann et al. (2018)). Furthermore, we used Ensembl Variant Effect Predictor (McLaren et al., 2016) to extract a list of genes affected by the top SNPs in these loci, of which multiple genes were previously reported to be associated with testosterone or the cholesterol level (Table 1).

**Table 1.**
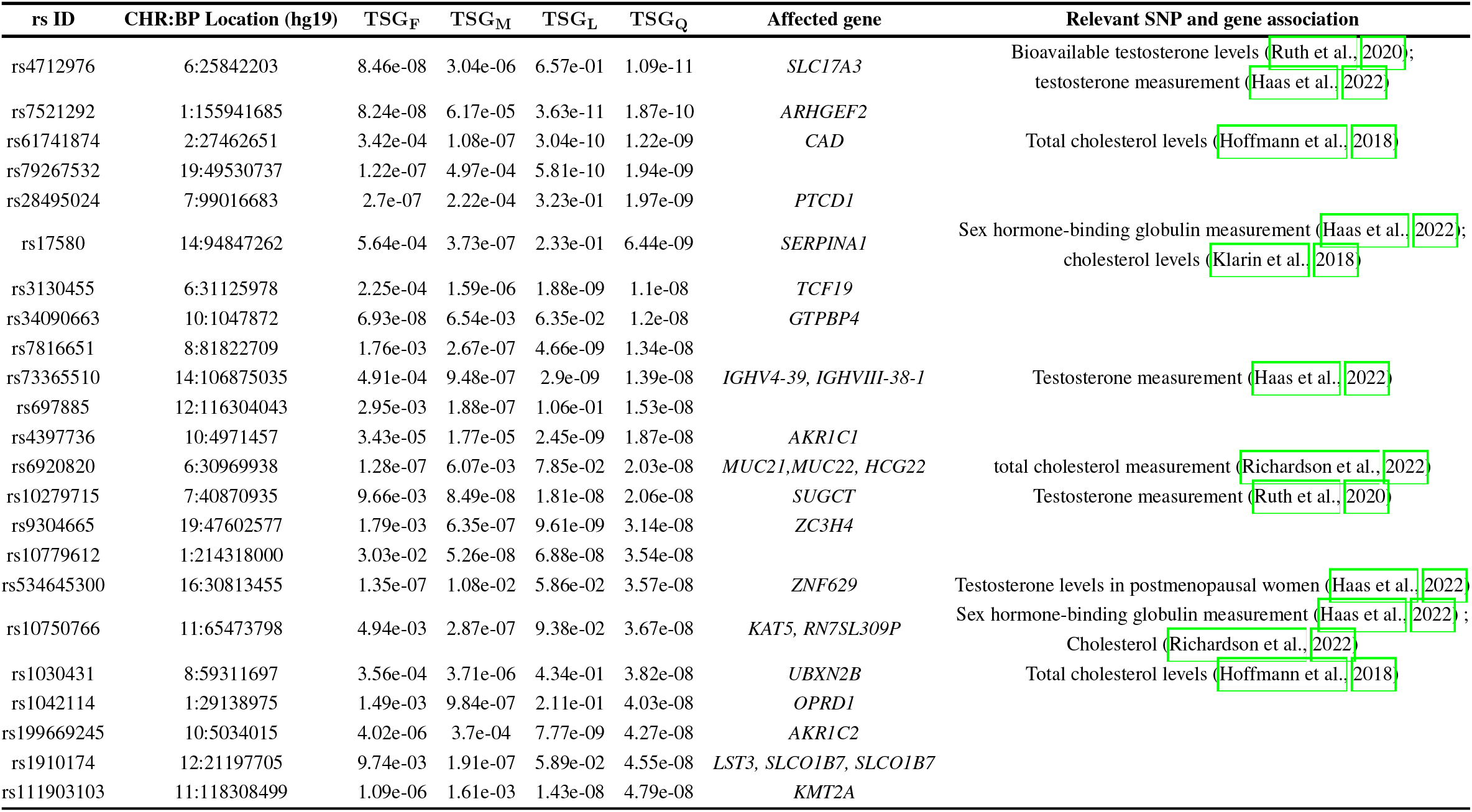
Evidence for biological relevance of the SNPs and genes in the loci uniquely identified by *TSG_Q_* in the UK Biobank data. Here *TSG_F_* and *TSG_M_* are sex-stratified association results with testosterone from the GWAS of Sinnott-Armstrong et al. (2021), which were also replicated by this study. The omnibus *TSG_Q_* results were obtained using equation 2, and the traditional meta-analysis *TSG_L_* results were obtained using equation 1. The p-values were calculated based on the summary statistics, *TSG_F_* and *TSG_M_*. Relevant SNP associations were based on previously reported association from GWAS catalog (Buniello et al., 2018); Affected genes and relevant gene association were extracted from Ensembl Variant Effect Predictor (McLaren et al., 2016).We ordered the rows based on *TSG_Q_* column and excluded four SNPs that could be annotated to the GWAS Catalog.

## 5 DISCUSSION

In this report, we reviewed three classes of association testing methods that account for genetic effect heterogeneity, namely the sex-stratified approach, sex-combined approaches based on summary statistics, and sex-combined approaches based on individual data. We conducted extensive simulation studies and an application study, leveraging the sex-stratified GWAS summary statistics of Sinnott-Armstrong et al. (2021) which studied the testosterone levels in the UK Biobank data, and we gained several insights.

First, our simulation results confirmed that there is no uniformly most powerful test in this study setting. Instead, the relative power between these tests will depend on the underlying genetic models, and may substantially change between different scenarios. For example, *T_main_* from the standard genetic main effect model is optimal when the true genetic effects are homogeneous between female and male. But, when the effects are in opposite directions with similar magnitudes, *T_main_* has no power to detect association. Conversely, the female-only sex-stratified test *TSG_F_* is optimal when the effect is only present in female, but it loses substantial power when the effect is also present in male.

Second, our simulation results revealed that, *T I_joint_*, the 2 df joint test of both genetic main effect and SNP×Sex interaction effect is most robust across all the scenarios considered. It remains powerful even if there was no SNP×Sex interaction or the effect was only present in one group. For each scenario considered, *T I_joint_* is either the most powerful test or only has a small amount of power loss as compared with the oracle method for that particular scenario, e.g. *T_main_* under the homogeneous genetic effect scenario A1, and *TSG_F_* under the female-only genetic effect scenario A2. The robustness of the joint test was previously noted, for example by Aschard et al. (2010): “In situations where the marginal test had better power than the joint test, the difference was small (less than 10% absolute power). Additionally, a recent work in a different study setting (Chen et al., 2021), comparing the standard 1 df additive test with the 2 df genotypic test, showed that “under additivity, the maximum power loss of a 2 df genotypic test is [theoretically] capped at 11.4% across all parameter values and sample sizes.” Indeed, in our study setting when *T I_joint_* was not the most powerful test, its maximum empirical power loss was 11.51% across all scenarios considered. In contrast, the maximum loss of power were 71.19% for *TSG_F_* and 96.96% for *T_main_*. Additionally, the power of *T I_joint_* dominates the minimum p-value approach *TSG_min_* in almost all cases considered, except when genetic effect is only present in female where *TSG_min_* is only slightly more powerful (at most 4.23% more) than *T I_joint_*.

Third, the power curves of *T I_joint_* (jointly testing SNP main and SNP×Sex interaction) and *TSG_Q_* (‘meta’ combining 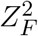 and 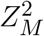) overlapped. This is not surprising given the earlier work of Aschard et al. (2010), which examined Score tests derived from the standard regression model where residual variance is not sex-specific. The empirical evidence we observed here deserves further investigation, and a theoretical result for the equivalence between *T I_joint_* and *TSG_Q_* would complement to the known result between *T_main_* (the traditional GWAS with no SNP×Sex interaction but including sex as a covariate) and *TSG_L_* (the traditional meta-analysis combining *Z_F_* and *Z_M_*) (Lin and Zeng, 2010), as well as the earlier work. The work of Aschard et al. (2010)

We emphasize that *T I_joint_* and *TSG_Q_* (if only sex-stratified summary statistics are available) are recommended for powerful association testing, but not for effect size estimation. Thus, *T I_joint_* and *TSG_Q_* could be applied as an screening approach for GWAS to identify significant variants. Subsequent refined analysis on these variants, such as the sex-stratified analysis, could be conducted to provide a complete picture of genetic effect heterogeneity.

As sex can be modelled statistically as an environmental factor, our observation here complements the work in Kraft et al. (2007), which focused on *T I_joint_* for gene×environmental interaction analysis and concluded that “the joint test [is] an attractive tool for large-scale association scans where the true gene×environment interaction model is unknown.” Indeed, even if the competing tests include the different data-integration methods such as the meta-analysis (*TSG_L_*), the minimal p-value approach (*TSG_min_*), and the omnibus test (*TSG_Q_*), *T I_joint_* remains competitive. This insight also challenges the current practice of analyzing samples from different populations separately. Instead, jointly analyzing all samples available and testing genetic main and gene-population effects simultaneously can be an useful alternative worth further investigation.

Finally, although we considered three different and complementing scenarios (A1, A2 and A3) for the true genetic model, simulation studies are never exhaustive, covering all possible alternatives. Indeed, when the residual distributions of phenotypic simulation model is non-normal and combined with the relatively small sample sizes, we observed inflated type I error at the extreme tail. Another future direction is to derive the non-centrality parameter of each test as a function of the minor allele frequency of a SNP, sex-specific effects of the SNP, sample size, male-to-female sample size ratio, as well as the random error. The analytical results will provide a complete picture for the power similarities and differences between the different tests.

## AUTHOR CONTRIBUTIONS

BL developed the method, performed the analyses, summarized the results, and drafted the manuscript. LS conceptualized and supervised the study. Both authors read and approved the final manuscript.

## ACKNOWLEDGMENTS

We thank Dr. Andrew Paterson, Dr. Shelley Bull and Dr. Lisa Strug for their helpful comments and discussion. BL is a trainee of the CANSSI-ONTARIO STAGE (Strategic Training for Advanced Genetic Epidemiology) training program at the University of Toronto.

