## Supplemental figures for "Better together against genetic effect heterogeneity: a sex-combined interaction analysis of testosterone levels in the UK Biobank data"

### Supplemental Material

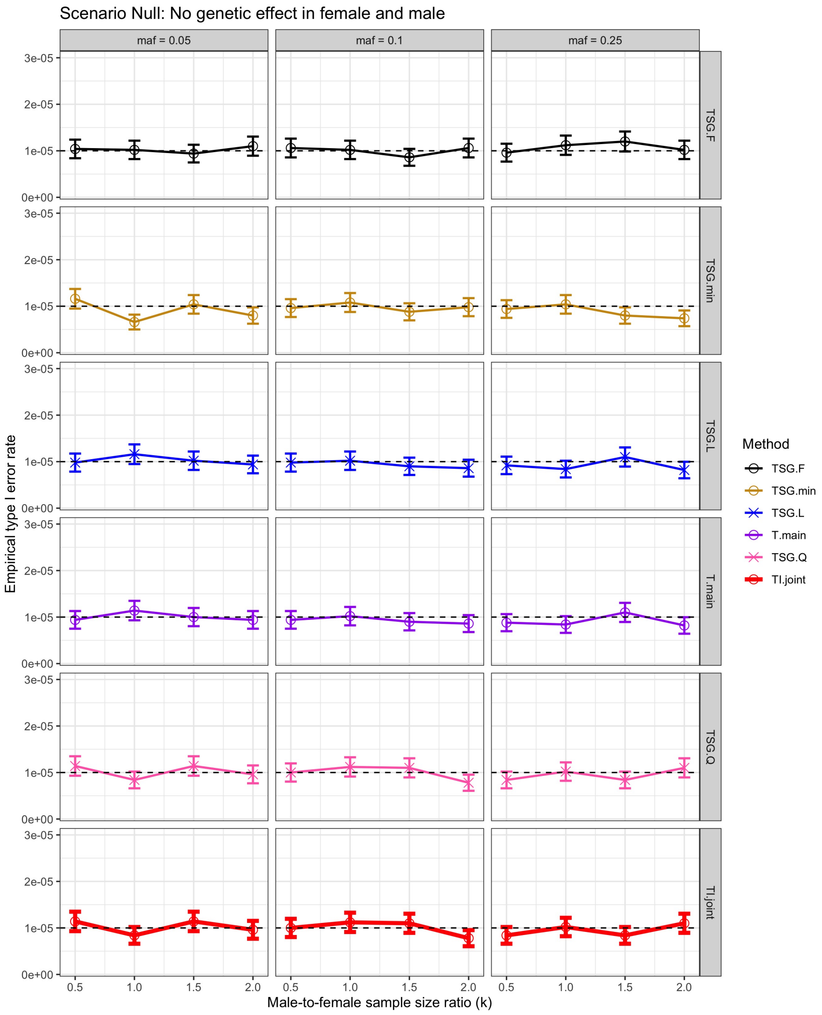


**Figure S1. The empirical type I error rates at the nominal $\alpha = 10^{-5}$ based on $R = 5 \times 10^6$ replications and their 95% confidence intervals under the null scenario, with $n_f = 5,000$, across different $k = n_m/n_f$ and $MAF$.** Six association testing methods were evaluated: (1) The female-only sex-stratified analysis ($TSG_F$); (2) The minimum p-value of female and male sex-stratified analysis ($TSG_{min}$); (3) A meta-analysis combining Z-scores ($TSG_L$); (4) The main-effect-only model ($T_{main}$); (5) An omnibus test combining squared Z-scores ($TSG_Q$), the recommended test when only sex-stratified summary statistics are available; (6) The recommended joint test of both the main and interaction effects ($TI_{joint}$). The dashed line indicates the nominal type I error rate of 1e-5. The 95% Binomial proportion confidence interval is constructed by formula: $\left(\hat{\alpha} - 1.96\sqrt{(1-\hat{\alpha})\hat{\alpha}/R}, \hat{\alpha} + 1.96\sqrt{(1-\hat{\alpha})\hat{\alpha}/R}\right)$, where $\hat{\alpha}$ is the empirical type I error rate.

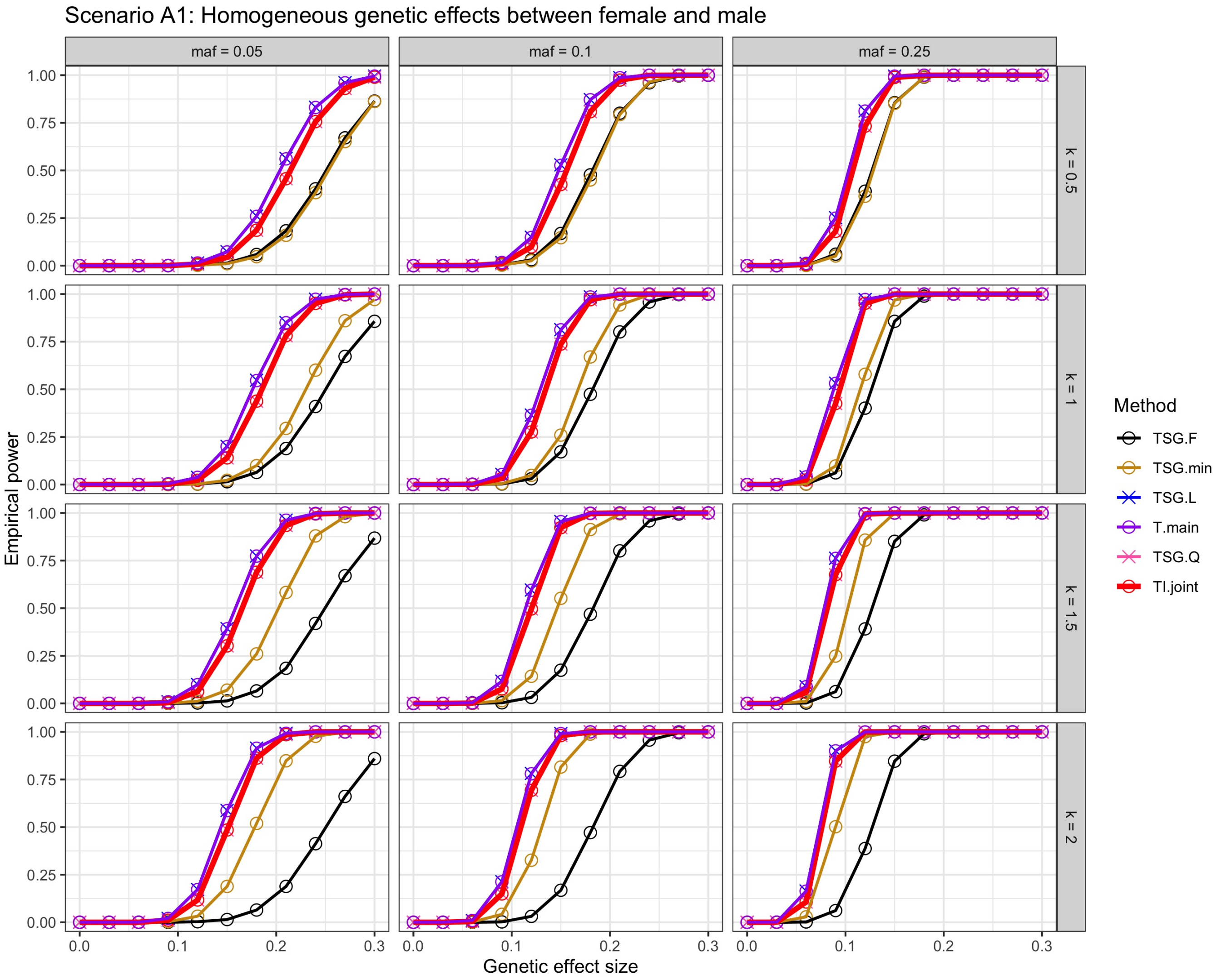


**Figure S2. Power comparison at $\alpha = $ 5e-8 with female sample sizes $n_f = 5,000$ under the alternative scenario A1 (homogeneous genetic effects between female and male), stratified by MAF and male-to-female sample size.** The genetic effect sizes ranged from 0 to 0.3. Six association testing methods were evaluated: (1) The female-only sex-stratified analysis ($TSG_F$); (2) The minimum p-value of sex-stratified analysis ($TSG_{min}$); (3) A meta-analysis combining Z-scores ($TSG_L$); (4) The main-effect-only model ($T_{main}$); (5) An omnibus test combining squared Z-scores ($TSG_Q$), the recommended test when only sex-stratified summary statistics are available; (6) The recommended joint test of both the main and interaction effects ($TI_{joint}$).

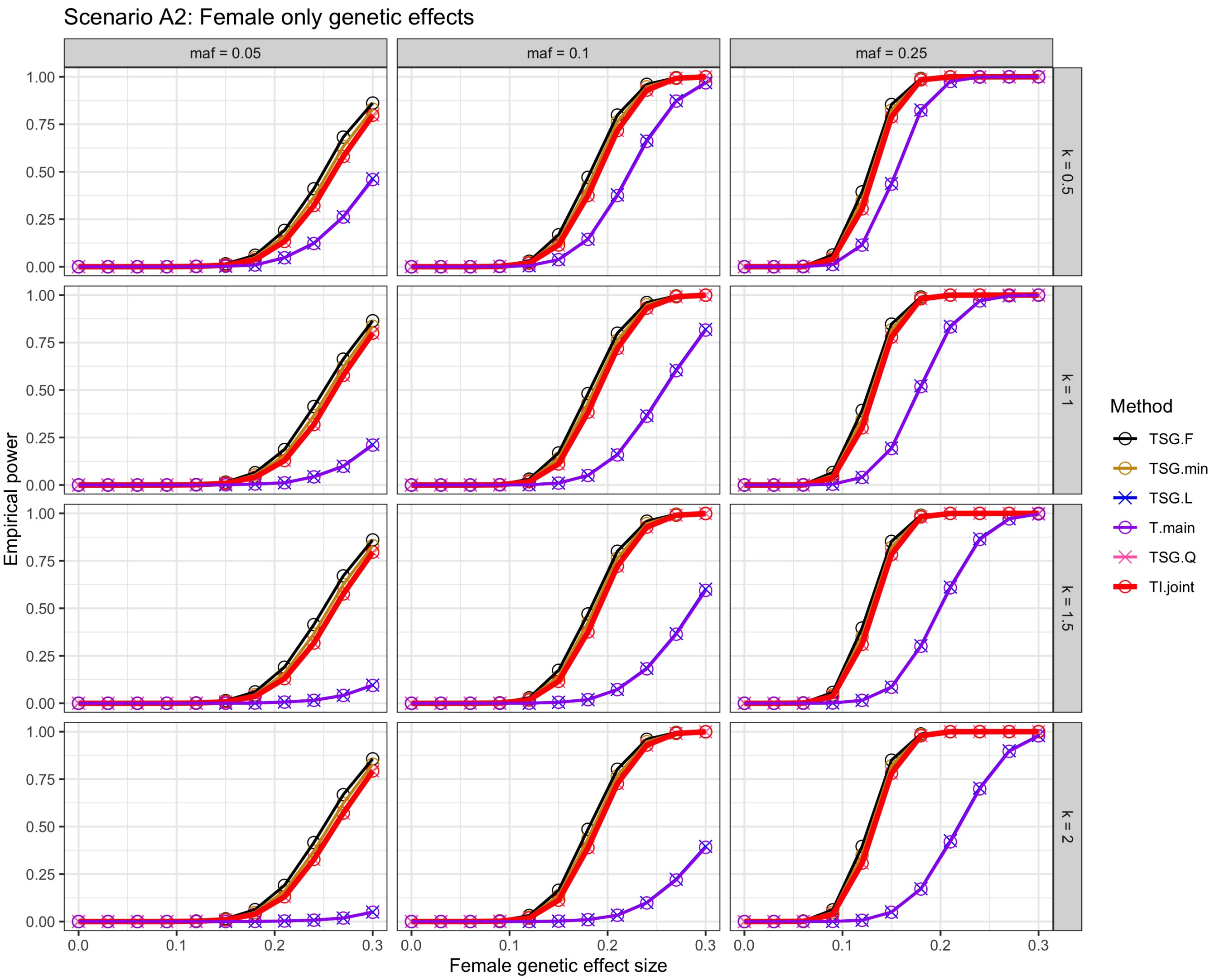


**Figure S3. Power comparison at $\alpha = 5\text{e-}8$ with female sample sizes $n_f = 5,000$ under the alternative scenario A2 (female only genetic effect), stratified by MAF and male-to-female sample size.** The genetic effect sizes ranged from 0 to 0.3. Six association testing methods were evaluated: (1) The female-only sex-stratified analysis ($TSG_F$); (2) The minimum p-value of sex-stratified analysis ($TSG_{min}$); (3) A meta-analysis combining Z-scores ($TSG_L$); (4) The main-effect-only model ($T_{main}$); (5) An omnibus test combining squared Z-scores ($TSG_Q$), the recommended test when only sex-stratified summary statistics are available; (6) The recommended joint test of both the main and interaction effects ($TI_{joint}$).

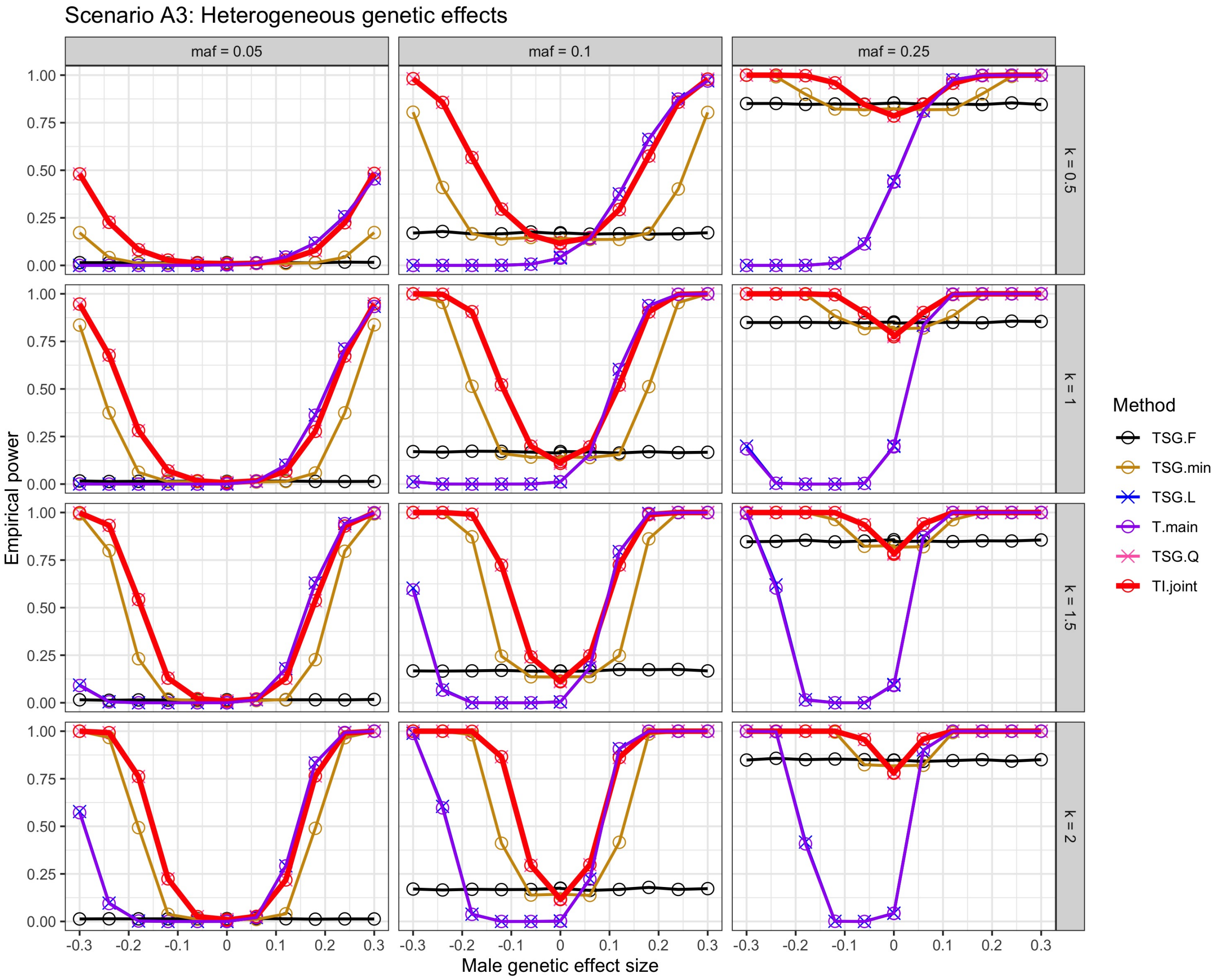


**Figure S4. Power comparison at $\alpha = $ 5e-8 with female sample sizes $n_f = 5,000$ under the alternative scenario A3 (heterogeneous effects), stratified by MAF and male-to-female sample size.** The genetic effect in female was kept at 0.15, while the effect in male ranged from -0.3 to 0.3. Six association testing methods were evaluated: (1) The female-only sex-stratified analysis ($TSG_F$); (2) The minimum p-value of sex-stratified analysis ($TSG_{min}$); (3) A meta-analysis combining Z-scores ($TSG_L$); (4) The main-effect-only model ($T_{main}$); (5) An omnibus test combining squared Z-scores ($TSG_Q$), the recommended test when only sex-stratified summary statistics are available; (6) The recommended joint test of both the main and interaction effects ($TI_{joint}$).

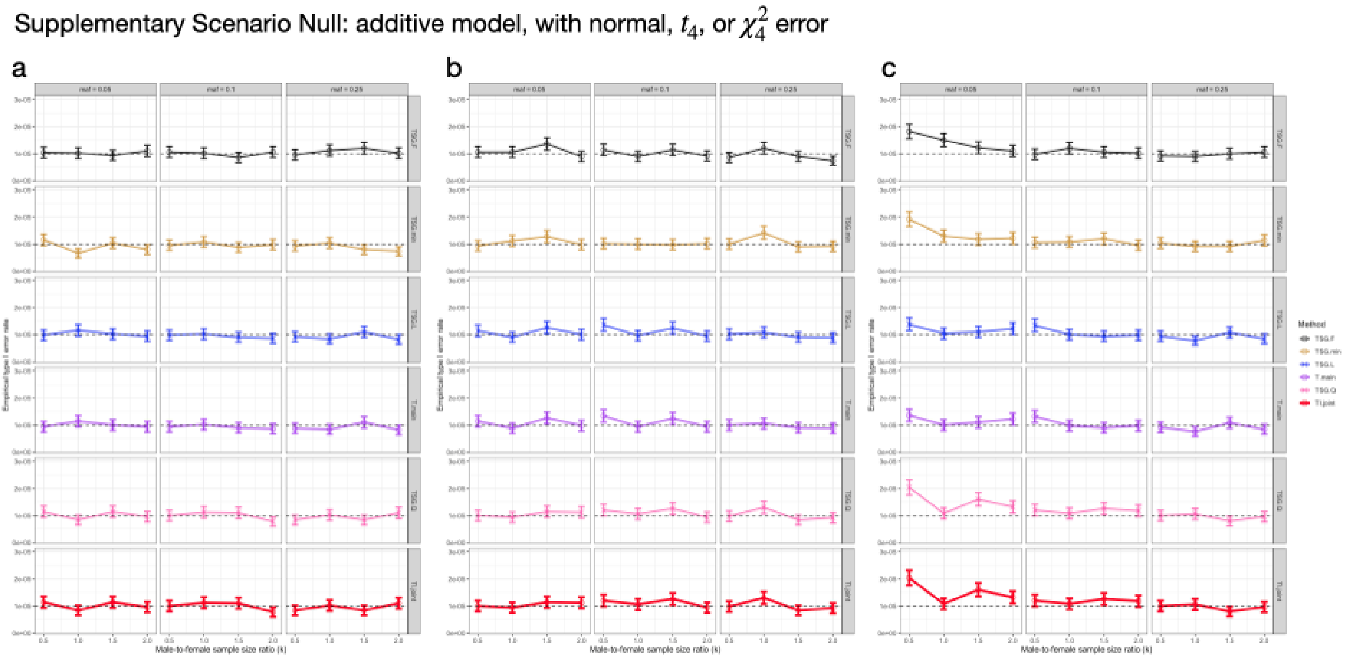


**Figure S5. Results of sensitivity study: non-normal residuals.** The empirical type I error rates at the nominal $\alpha = $ 1e-5 based on $R = 5 \times 10^6$ replications and their 95% confidence intervals under the null scenario, with $n_f = n_m = 5{,}000$, stratified by residual distributions (a) standard normal, (b) $t_4$ and (c) $\chi^2_4$. The genotypes simulated under additive model. Six association testing methods were evaluated: (1) The female-only sex-stratified analysis ($TSG_F$); (2) The minimum p-value of sex-stratified analysis ($TSG_{min}$); (3) A meta-analysis combining Z-scores ($TSG_L$); (4) The main-effect-only model ($T_{main}$); (5) An omnibus test combining squared Z-scores ($TSG_Q$), the recommended test when only sex-stratified summary statistics are available; (6) The recommended joint test of both the main and interaction effects ($TI_{joint}$). The dashed line indicates the nominal type I error rate of 1e-5. The 95% Binomial proportion confidence interval is constructed by formula: $\left(\hat{\alpha} - 1.96\sqrt{\frac{(1-\hat{\alpha})\hat{\alpha}}{R}}, \hat{\alpha} + 1.96\sqrt{\frac{(1-\hat{\alpha})\hat{\alpha}}{R}}\right)$, where $\hat{\alpha}$ is the empirical type I error rate.

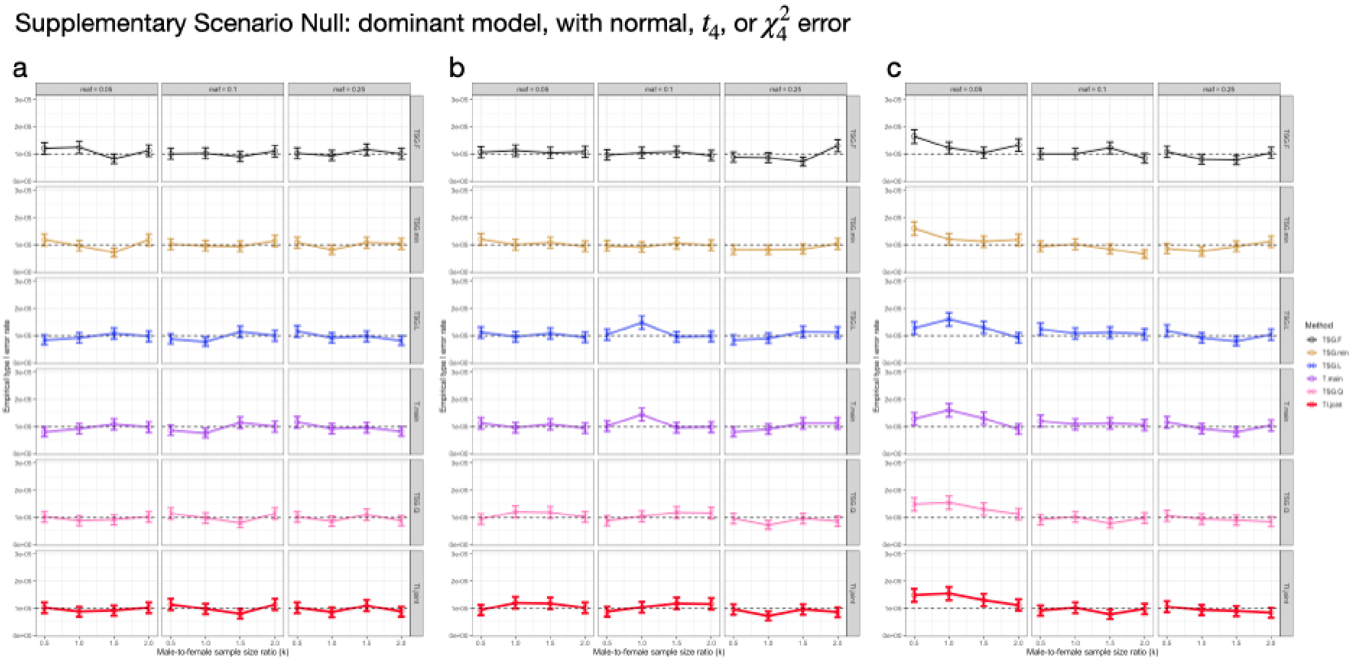


**Figure S6. Results of sensitivity study: non-additive generating model**. The empirical type I error rates at the nominal $\alpha = $ 1e-5 based on $R = 5 \times 10^6$ replications and their 95% confidence intervals under the null scenario, with $n_f = n_m = 5,000$, stratified by residual distributions (a) standard normal, (b) $t_4$ and (c) $\chi^2_4$. The genotypes simulated under dominant model. Six association testing methods were evaluated: (1) The female-only sex-stratified analysis ($TSG_F$); (2) The minimum p-value of sex-stratified analysis ($TSG_{min}$); (3) A meta-analysis combining Z-scores ($TSG_L$); (4) The main-effect-only model ($T_{main}$); (5) An omnibus test combining squared Z-scores ($TSG_Q$), the recommended test when only sex-stratified summary statistics are available; (6) The recommended joint test of both the main and interaction effects ($TI_{joint}$). The dashed line indicates the nominal type I error rate at $10^{-5}$. The 95% Binomial proportion confidence interval is constructed by formula: $\left(\hat{\alpha} - 1.96\sqrt{\frac{(1-\hat{\alpha})\hat{\alpha}}{R}}, \hat{\alpha} + 1.96\sqrt{\frac{(1-\hat{\alpha})\hat{\alpha}}{R}}\right)$, where $\hat{\alpha}$ is the empirical type I error rate.

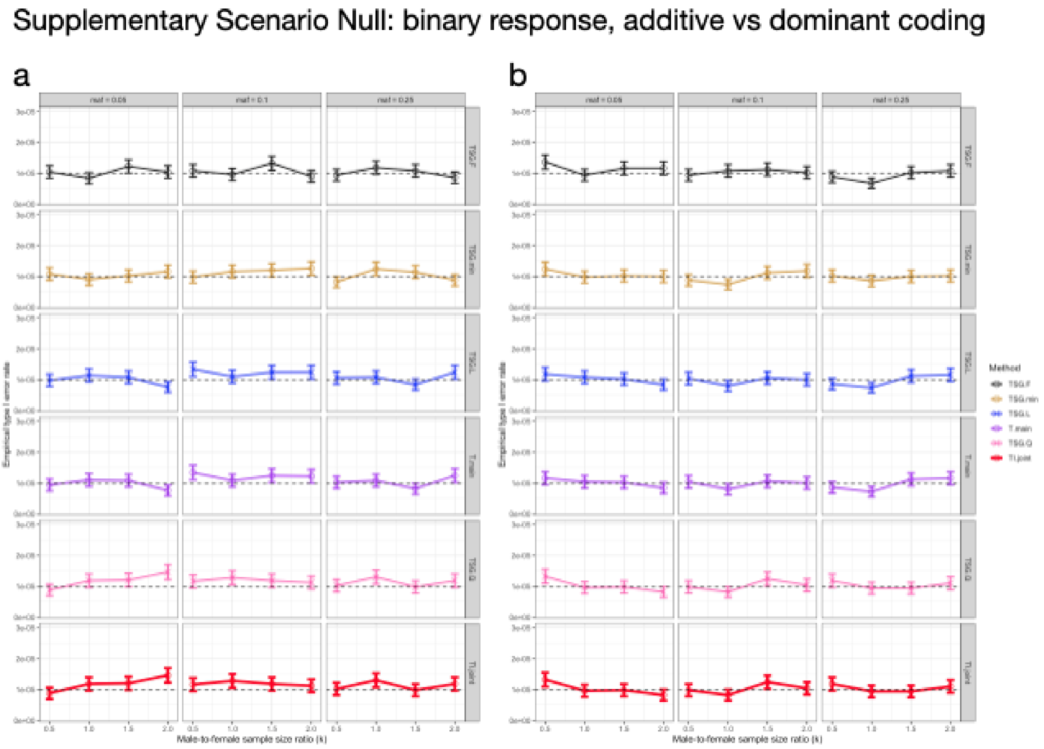


**Figure S7. Results of sensitivity study: binary trait and non-additive generating model.** The empirical type I error rates with binary response at the nominal $\alpha$ = 1e-5 based on $R = 5 \times 10^6$ replications and their 95% confidence intervals under the null scenario, with $n_f = n_m = 5,000$. We simulated data from (a) additive and (b) dominant genetic models respectively, although we assume additive model for association testing. Six association testing methods were evaluated: (1) The female-only sex-stratified analysis ($TSG_F$); (2) The minimum p-value of sex-stratified analysis ($TSG_{min}$); (3) A meta-analysis combining Z-scores ($TSG_L$); (4) The main-effect-only model ($T_{main}$); (5) An omnibus test combining squared Z-scores ($TSG_Q$), the recommended test when only sex-stratified summary statistics are available; (6) The recommended joint test of both the main and interaction effects ($TI_{joint}$). The dashed line indicates the nominal type I error rate at $10^{-5}$. The 95% Binomial proportion confidence interval is constructed by formula: $\left(\hat{\alpha} - 1.96\sqrt{\frac{(1-\hat{\alpha})\hat{\alpha}}{R}}, \hat{\alpha} + 1.96\sqrt{\frac{(1-\hat{\alpha})\hat{\alpha}}{R}}\right)$, where $\hat{\alpha}$ is the empirical type I error rate.

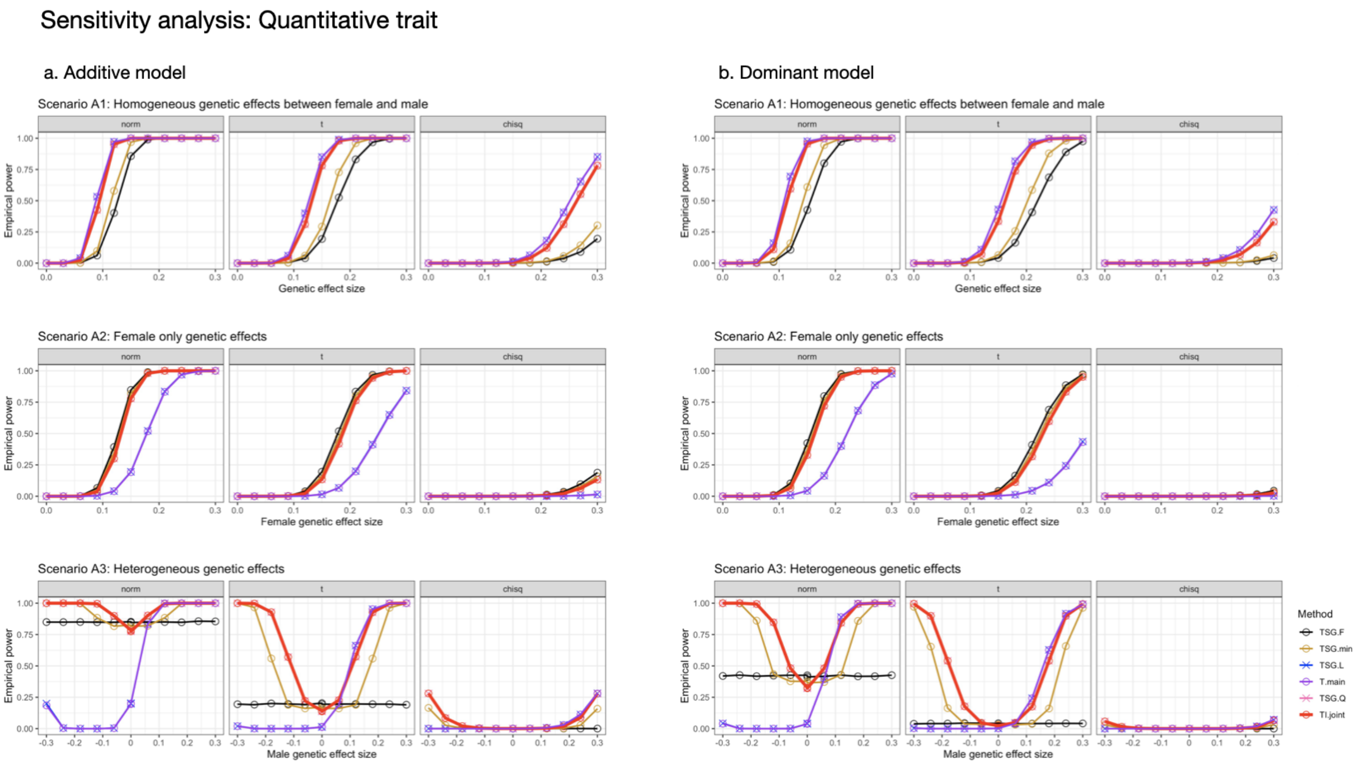


**Figure S8. Quantitative trait power at $\alpha$ = 5e-8 under the three alternative scenarios, stratified by error distributions and genetic models.** Sample sizes $n_f = n_m = 5,000$, and $MAF = 0.25$. The columns correspond to the additive and dominant genetic models respectively, although we assume additive model for association testing. Six association testing methods were evaluated: (1) The female-only sex-stratified analysis ($TSG_F$); (2) The minimum p-value of sex-stratified analysis ($TSG_{min}$); (3) A meta-analysis combining Z-scores ($TSG_L$); (4) The main-effect-only model ($T_{main}$); (5) An omnibus test combining squared Z-scores ($TSG_Q$), the recommended test when only sex-stratified summary statistics are available; (6) The recommended joint test of both the main and interaction effects ($TI_{joint}$).

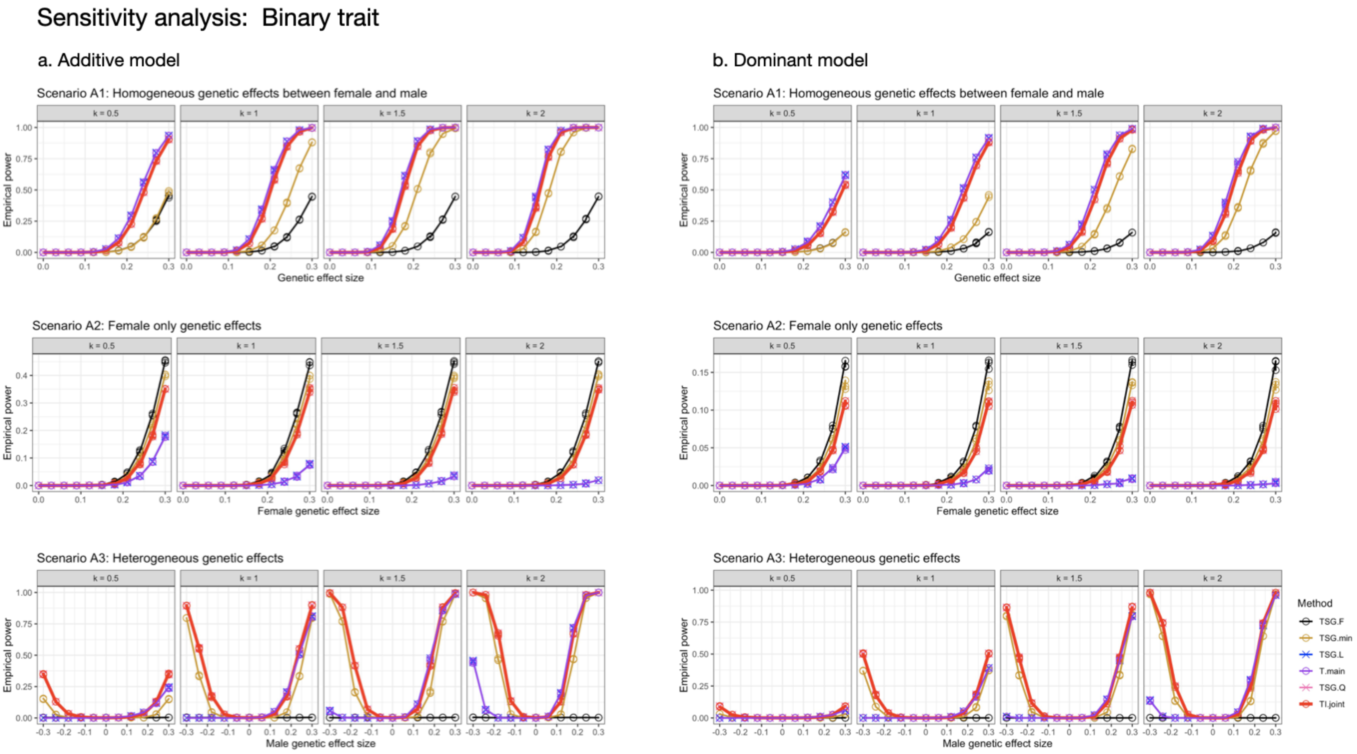


**Figure S9. Binary trait power at $\alpha = $ 5e-8 under the three alternative scenarios, stratified by genetic models and female-to-male sample size ratio (k).** Sample sizes $n_f = 5,000$, $n_m = n_f \times k$, and $MAF = 0.25$. The columns correspond to the additive and dominant genetic models respectively, although we assume additive model for association testing. Six association testing methods were evaluated: (1) The female-only sex-stratified analysis ($TSG_F$); (2) The minimum p-value of sex-stratified analysis ($TSG_{min}$); (3) A meta-analysis combining Z-scores ($TSG_L$); (4) The main-effect-only model ($T_{main}$); (5) An omnibus test combining squared Z-scores ($TSG_Q$), the recommended test when only sex-stratified summary statistics are available; (6) The recommended joint test of both the main and interaction effects ($TI_{joint}$).

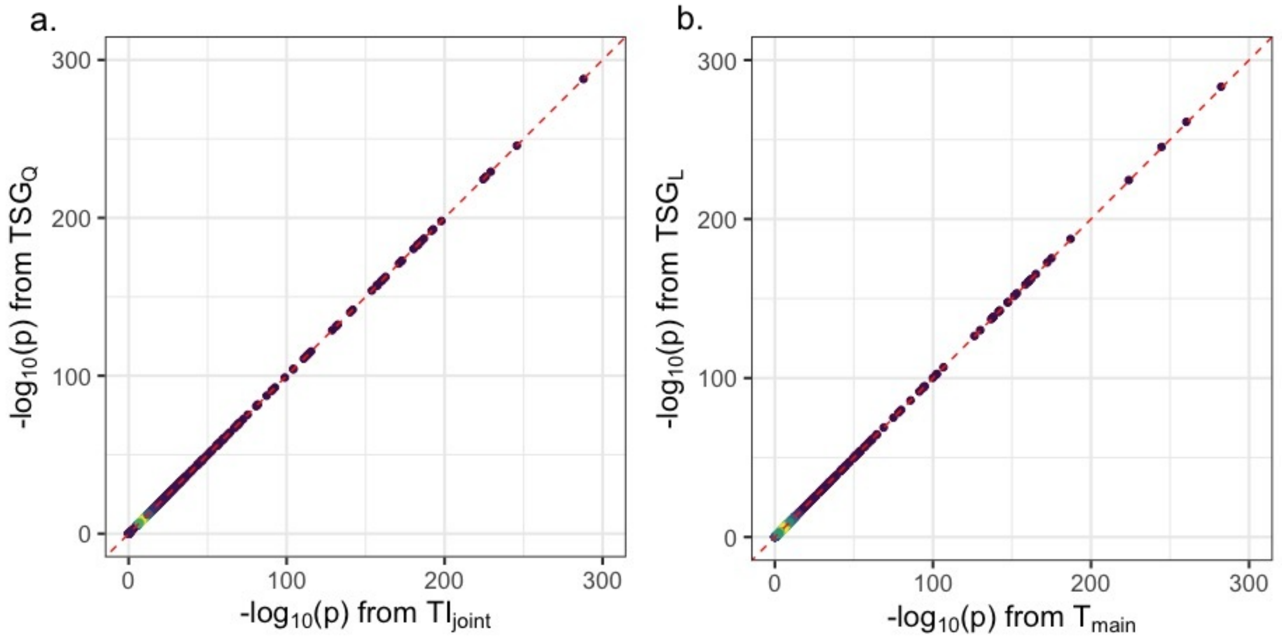


**Figure S10. PP plots, from the UKB application, shows the equivalence between (a) $TSG_Q$ vs $TI_{joint}$ and (b) $TSG_L$ and $T_{main}$.** Here quadratic omnibus test $TSG_Q$ and traditional meta-analysis $TSG_L$ were obtained using equation (2) and (1). $TI_{joint}$ is the 2 d.f. main and interaction testing approach, and $T_{main}$ is the main-effect only testing approach (and adjusting for sex effect). Note that both $TI_{joint}$ and $T_{main}$ were derived from regression models with sex-specific residual variances.

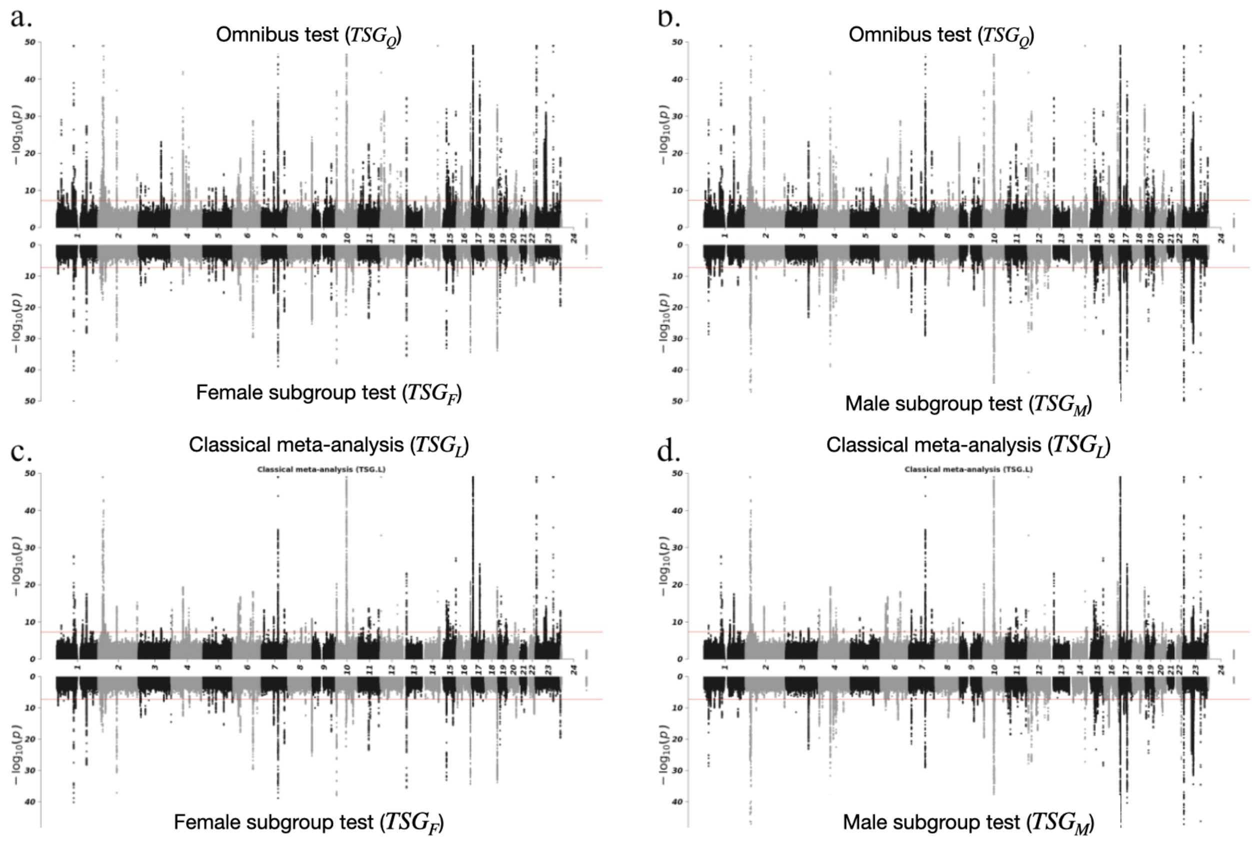


**Figure S11. Miami plots with the maximum $-\log_{10} p$ is truncated at 50 for the UKB application,** comparing the $-\log_{10}(p)$ between tests based on (a.) $TSG_Q$ and $TSG_F$, (b.) $TSG_Q$ and $TSG_M$, (c.) $TSG_L$ and $TSG_F$, (d.) $TSG_L$ and $TSG_M$ of GWAS on testosterone level from Sinnott-Armstrong et al. (2021). Here $TSG_F$, $TSG_M$ are the test statistics of female- and male-only subgroup analysis respectively. The omnibus test combining squared Z-scores ($TSG_Q$) and meta-analysis combining Z-scores ($TSG_L$) were calculated by combining $TSG_F$ and $TSG_M$. The red line indicates genome-wide significant threshold at $5 \times 10^{-8}$.

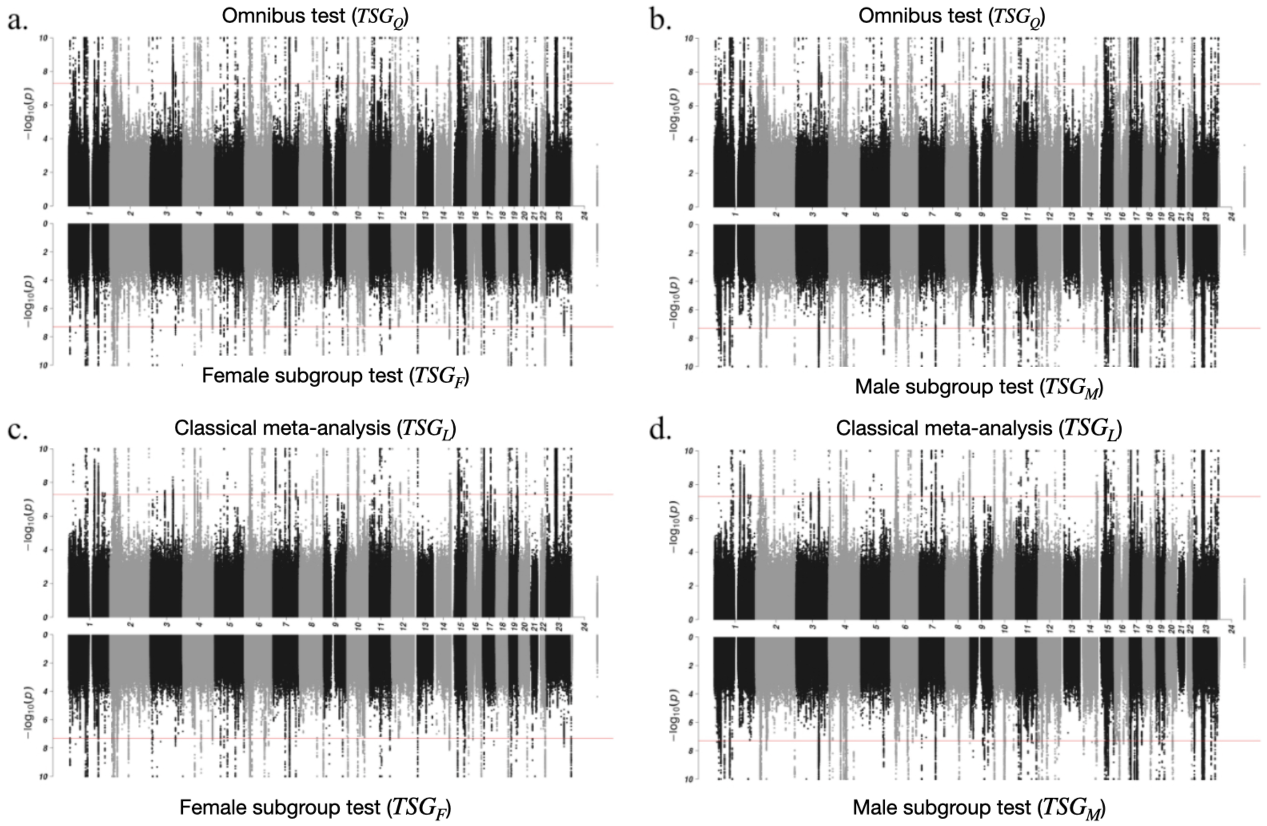


**Figure S12. Miami plots with the maximum $-\log_{10} p$ is truncated at 10 for the UKB application,** comparing the $-\log_{10}(p)$ between tests based on (a.) $TSG_Q$ and $TSG_F$, (b.) $TSG_Q$ and $TSG_M$, (c.) $TSG_L$ and $TSG_F$, (d.) $TSG_L$ and $TSG_M$ of GWAS on testosterone level from Sinnott-Armstrong et al. (2021). Here $TSG_F$, $TSG_M$ are the test statistics of female- and male-only subgroup analysis respectively. The omnibus test combining squared Z-scores ($TSG_Q$) and meta-analysis combining Z-scores ($TSG_L$) were calculated by combining $TSG_F$ and $TSG_M$. The red line indicates genome-wide significant threshold at $5 \times 10^{-8}$.

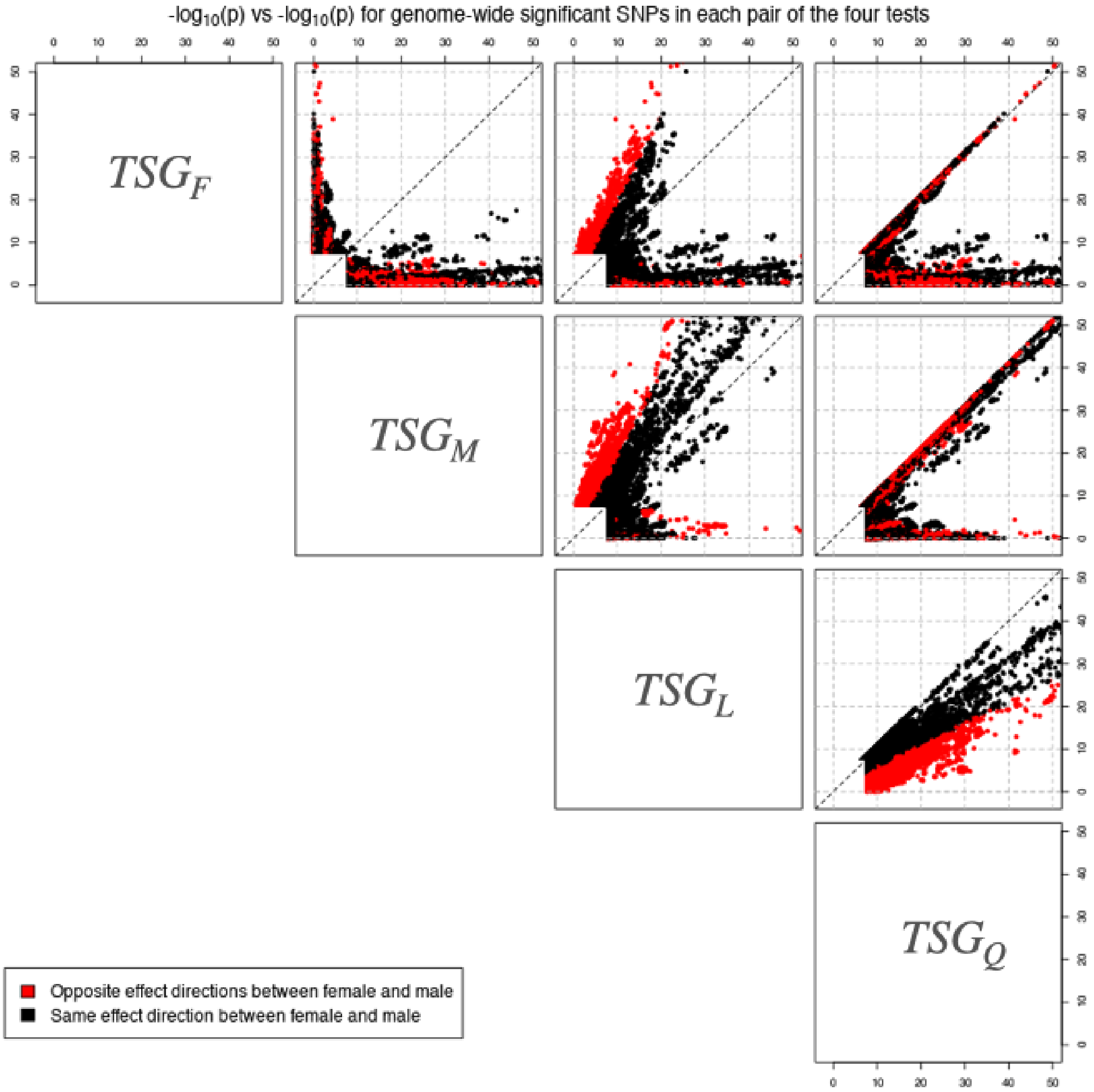


**Figure S13. Pairwise PP plots from the UKB application, with p-values truncated at 50 for better visualization,** comparing the $-\log_{10} p$-values between the female-only subgroup analysis ($TSG_F$), male-only subgroup analysis ($TSG_M$), the classical meta-analysis ($TSG_L$), and the omnibus test combining squared Z-scores ($TSG_Q$). In each sub-plot, the points represent the variants reported genome-wide significant, at 5e-8, in *any* of the two corresponding testing methods. The p-values for $TSG_F$ and $TSG_M$ were obtained (and replicated) from the summary statistics of Sinnott-Armstrong et al. (2021), while $TSG_Q$ and $TSG_L$ were calculated by combining $TSG_F$ and $TSG_M$.

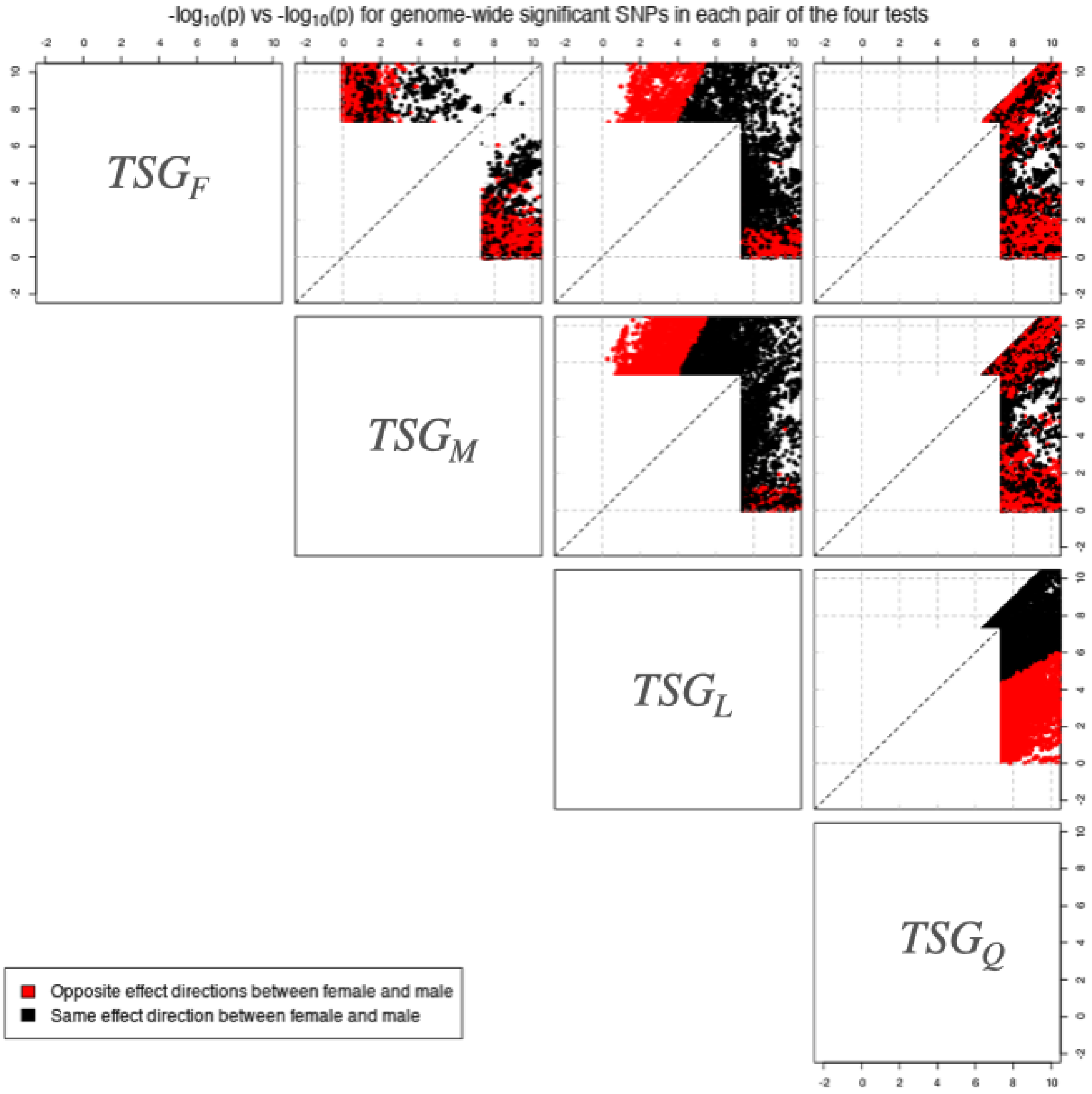


**Figure S14. Pairwise PP plots from the UKB application, with p-values truncated at 10 for better visualization,** comparing the $-\log_{10} p$-values between the female-only subgroup analysis ($TSG_F$), male-only subgroup analysis ($TSG_M$), the classical meta-analysis ($TSG_L$), and the omnibus test combining squared Z-scores ($TSG_Q$). In each sub-plot, the points represent the variants reported genome-wide significant, at 5e-8, in *any* of the two corresponding testing methods. The p-values for $TSG_F$ and $TSG_M$ were obtained (and replicated) from the summary statistics of Sinnott-Armstrong et al. (2021), while $TSG_Q$ and $TSG_L$ were calculated by combining $TSG_F$ and $TSG_M$.
